# Individually Unique Dynamics of Cortical Connectivity Reflect the Ongoing Intensity of Chronic Pain

**DOI:** 10.1101/2021.06.30.450553

**Authors:** Astrid Mayr, Pauline Jahn, Bettina Deak, Anne Stankewitz, Vasudev Devulapally, Viktor Witkovsky, Olaf Dietrich, Enrico Schulz

## Abstract

**Background:** Chronic pain diseases are characterised by an ongoing and fluctuating endogenous pain, yet it remains to be elucidated how this is reflected by the dynamics of ongoing functional cortical connections. The present study addresses this disparity by taking the individual perspective of pain patients into account, which is the varying intensity of endogenous pain.

**Methods:** To this end, we investigated the cortical encoding of 20 chronic back pain patients and 20 chronic migraineurs in four repeated fMRI sessions. During the recording, the patients were asked to continuously rate their pain intensity. A brain parcellation approach subdivided the whole brain into 408 regions. A 10 s sliding-window connectivity analysis computed the pair-wise and time-varying connectivity between all brain regions across the entire recording period. Linear mixed effects models were fitted for each pair of brain regions to explore the relationship between cortical connectivity and the observed trajectory of the patients’ fluctuating endogenous pain.

**Results:** Two pain processing entities were taken into account: pain intensity (high, middle, low pain) and the direction of pain intensity changes (rising vs. falling pain). Overall, we found that periods of high and increasing pain were predominantly related to low cortical connectivity. For chronic back pain this applies to the pain intensity-related connectivity for limbic and cingulate areas, and for the precuneus. The change of pain intensity was subserved by connections in left parietal opercular regions, right insular regions, as well as large parts of the parietal, cingular and motor cortices. The change of pain intensity direction in chronic migraine was reflected by decreasing connectivity between the anterior insular cortex and orbitofrontal areas, as well as between the PCC and frontal and ACC regions.

**Conclusions:** Interestingly, the group results were not mirrored by the individual patterns of pain-related connectivity, which is suggested to deny the idea of a common neuronal core problem for chronic pain diseases. In a similar vein, our findings are supported by the experience of clinicians, who encounter patients with a unique composition of characteristics: personality traits, various combinations of symptoms, and a wide range of individual responses to treatment. The diversity of the individual cortical signatures of chronic pain encoding results adds to the understanding of chronic pain as a complex and multifaceted disease. The present findings support recent developments for more personalised medicine.

## Introduction

Functional connectivity (FC) in the brain is a measure of spatio-temporal correlations of fluctuating time-series of distinct regions. It has become essential for exploring the functional network structure of both healthy and diseased brains ^1–4^ as alterations in functional connectivity patterns due to neurological or psychiatric disorders, as well as changes due to long-lasting chronic pain conditions, have been reported ^5–7^. Functional connectivity in chronic pain conditions have been studied by analysing intrinsic cortical networks as well as by computing seed-based connections of one or more predefined brain regions.

A number of studies have found changes in intrinsic network connectivity across several chronic pain disorders, particularly for the default mode network (DMN). Compared to healthy subjects, diabetic neuropathic pain patients show greater connectivity between the DMN and the anterior cingulate cortex (ACC) ^8^, and fibromyalgia patients show greater connectivity between the DMN and insular cortices ^9^. For chronic migraineurs (CM), disrupted functional connectivity between the DMN and the executive control network, but increased functional connectivity between the DMN and the left dorsal attention system, has been reported ^10^. Chronic back pain (CBP) patients were found to exhibit a general reorganisation of the DMN, with increased connectivity of the DMN to the medial prefrontal cortex (mPFC), ACC and left anterior insula, and decreased connectivity of mPFC with the precuneus ^11,12^.

Specifically for CM patients compared to episodic migraineurs (EM), a further study revealed greater connectivity within an intrinsic network consisting of the ACC, the anterior insula, the thalamus, the dorsolateral prefrontal cortex (DLPFC), the precuneus, the supramarginal gyrus, and the cerebellum. In addition, a greater connectivity between this network, the hypothalamus, and the dorsal raphe nuclei has been found ^13^. A further study reported decreased connectivity between the executive control network and the left dorsal attention system in CM compared to healthy subjects ^10^.

In a previous and complex whole-brain connectivity study using cortical parcellation, multiple changes for CBP were found, particularly for connections involving the frontal cortex and the ACC ^14^. Furthermore, CM patients showed stronger connectivity of the amygdalae with regions in the inferior temporal, prefrontal, cingulate, as well as the pre-and postcentral cortices compared to EM ^15^. Weaker connectivity was found between the right amygdala and occipital regions in CM compared to healthy subjects. Similar results were obtained by ^16^, who were identifying atypical resting-state functional connectivity e.g. of the anterior insula, amygdala, thalamus and the periaqueductal grey (PAG) in CM compared to controls. In addition, disease duration correlated with the strength of connectivity between the anterior insula, thalamic nuclei, and the PAG. CM patients were also found to show decreased connectivity within a central executive network between the right ventrolateral prefrontal cortex (PFC) and the thalamus, as well as between the left dorsal PFC and the dorsomedial PFC compared to controls ^17^. A further study investigated CBP patients in high and low pain conditions. High back pain intensity was associated with stronger FC of the primary somatosensory (S1) and motor (M1) cortices and the left superior frontal cortex ^18^. Moreover, CBP has been found to alter the connectivity between primary sensory networks compared to controls ^19^. Increased connectivity between the primary visual network and S1 was reported, and connectivity strength was further negatively correlated with the disease duration.

There are a number of challenges that hamper the formation of a unified framework for the understanding of the cortical underpinnings of chronic pain disease and may explain the often contradicting findings. *First*, the cause of the common low-frequency fluctuation of BOLD activity, which gives rise to distinct intrinsic networks, is not yet entirely understood. The BOLD fluctuations, consisting of peaks and troughs, may be related to the continuously-changing pain intensity. This relationship has not been investigated so far and network fluctuations that are unrelated to the ongoing experience of pain would be difficult to interpret. *Second*, many studies in the neuroimaging literature lack reliable cortical data with repeated measurements for reproducible and longitudinal observations ^6,20^, which may explain the heterogeneity of the DMN findings. Therefore, extended and repeated recordings are required to disentangle a subject’s stable signature from random momentary fluctuations of brain connectivity.

Here, we aimed to capture the individual endogenous pain experience for each subject. We utilised a whole-brain functional connectivity approach to identify coherent brain connectivity patterns of chronic pain encoding (CBP and CM). To this end, pain ratings were related to simultaneously obtained cortical data to characterise the fluctuating neuronal connectivity patterns that reflect the experience of chronic pain for each individual across several recordings. Multiple recordings for each individual enabled us to explore how well the results of group statistics reflect the individual processing of chronic pain.

## Material and Methods

### Participants

The study included 20 patients diagnosed with chronic back pain (CBP - 16 female; aged 44±13 years) and 20 patients with chronic migraine (CM - 18 female; aged 34±13 years). All participants gave written informed consent. The study was approved by the Ethics Committee of the Medical Department of the Ludwig-Maximilians-Universität München and conducted in conformity with the Declaration of Helsinki.

CBP patients were diagnosed according to the IASP criteria (The International Association for the Study of Pain) ^21^, which includes a disease duration of more than 6 months (mean CBP: 10±7 years). All patients were seen in a specialised pain unit. CM patients were diagnosed according to the ICHD-3 ^22^, defined as a headache occurring on 15 or more days/month for more than 3 months, which, on at least 8 days/month, has the features of migraine headache (mean CM: 15±12 years). All CM patients were seen in a tertiary headache centre.

All patients were permitted to continue their pharmacological treatment at a stable dose (Supplementary Tables 1 and 2). The patients did not report any other neurological or psychiatric disorders, or had contraindications for an MRI examination. Patients who had any additional pain were excluded. For all patients, pain was fluctuating and not constant at the same intensity level. Patients with no pain or headache attacks on the day of the measurement were asked to return on a different day. Patients were characterised using the German Pain Questionnaire (Deutscher Schmerzfragebogen) ^23^ and the German version of the Pain Catastrophizing Scale (PCS; Supplementary Tables 1 and 2) ^24^. The pain intensity describes the average pain in the last 4 weeks from 0 to 10 with 0 representing no pain and 10 indicating maximum imaginable pain (please note that this scale differs from the one used in the fMRI experiment). The German version of the Depression, Anxiety and Stress Scale (DASS) was used to rate depressive, anxiety, and stress symptoms over the past week ^25^.

Patients were compensated with 60€ for each session. In total nine screened patients were excluded: two patients developed additional pain during the study, the pain ratings of five patients were constantly increasing or decreasing throughout the pain rating experiment, and two patients were unable to comply with study requests. 36 patients were recorded four times across 6 weeks with a gap of at least 2 days (CBP = 9±12 days, CM = 12±19 days) between sessions. Four patients (2 CBP and 2 CM) were recorded three times. The additional pain in two patients would confound the analysis of the cortical processing of chronic pain. Furthermore, a steadily increasing time course of pain ratings is not suitable for fMRI analyses. The exclusion is an important prerequisite for the statistical independence of the three entities that describe the processing of chronic pain (see below).

### Experimental procedure

During the recording of fMRI, patients rated the intensity of their ongoing pain for 25 minutes using an MRI-compatible potentiometer slider ^26^. The scale ranged from 0 to 100 in steps of five with 0 representing no pain and 100 representing the highest experienced pain. On a light grey screen a moving red cursor on a dark grey bar (visual analogue scale, VAS) and a number above the bar (numeric analogue scale, NAS) were shown during the entire functional MRI session. The screen was visible through a mirror mounted on top of the MRI head coil. Patients were asked to look only at the screen and to focus on their pain. The intensity and the changes of perceived pain had to be indicated as quickly and accurately as possible. To minimise head movement, foams were placed around the head and patients were told to lie as still as possible.

### Data Acquisition

Data were recorded on a clinical 3T MRI scanner (Siemens Magnetom Skyra, Germany) using a 64-channel head coil. A T2*-weighted BOLD (blood oxygenation level dependent) gradient echo sequence with echo-planar image acquisition and a multiband factor of 2 was used with the following parameters: number of slices = 46; repetition time/echo time = 1550/30 ms; flip angle = 71°; slice thickness = 3 mm; voxel size = 3×3×3 mm^3^; field of view = 210 mm. 1000 volumes were recorded in 1550 seconds. Field maps were acquired in each session to control for B0-effects. For each patient, T1- and T2-weighted anatomical MRI images were acquired using the following parameters for T1: repetition time/echo time = 2060/2.17 ms; flip angle = 12°; number of slices = 256; slice thickness = 0.75 mm; field of view = 240 mm, and for T2: repetition time/echo time = 3200/560 ms; flip angle = 120°; number of slices = 256; slice thickness = 0.75 mm; field of view = 240 mm.

### Data processing - behavioural data

The rating data were continuously recorded with a variable sampling rate but downsampled offline at 10 Hz. To remove the same filtering effects from the behavioural data as from the imaging data, we applied a 400 s high-pass filter (see below). The rating data was convolved with a hemodynamic response function (HRF) implemented in SPM12 ^27^ with the following parameters: HRF = spm_hrf(0.1,[6 16 1 1 100 0 32]). The post-stimulus undershoot was minimised by the ratio of response to undershoot and motivated by the continuous and event-free fMRI design. For the statistical analysis, the resulting filtered time course was transferred to Matlab (Mathworks, USA; version R2018a) and downsampled to the sampling frequency of the imaging data (1/1.55 Hz).

To disentangle the distinct aspects of pain intensity (AMP - amplitude) from cortical processes related to the sensing of rising and falling pain, we computed the ongoing rate of change in the pain ratings (illustrated in Figure 1). The rate of change is calculated as the slope of the regression of the least squares line across a 3 s time window of the 10 Hz pain rating data (SLP - slope, encoded as 1, -1, and 0). Periods coded as 0 indicate time frames of constant pain. The absolute slope of pain ratings (aSLP - absolute slope, encoded as 0 and 1) represents periods of motor-related connectivity (slider movement), changes of visual input (each slider movement changes the screen), and decision-making (each slider movement prerequisites a decision to move). Periods coded as 0 indicate time frames of constant pain without the need to move the slider. We deliberately kept the SLP and aSLP as nominal variables; a higher velocity of slider movement or a faster change of pain intensity are unlikely to cause a proportional change in brain connectivity. The low correlations of the three entities (AMP, SLP, aSLP) indicate the independence of the vectors. The mean (±standard deviation) correlation coefficients (Fisher-z transformed) for all CBP/CM subjects for each of the variable pairs were: r(AMP, SLP) = 0.007(±0.04) / 0.02(±0.04); r(SLP, aSLP) = 0.06(±0.2) / 0.01(±0.12); r(AMP, aSLP) = 0.002(±0.08) / -0.004(±0.09). The rating time courses were required to fluctuate at a relatively constant level, in order to mitigate potential effects of order (e.g. in case of continuously rising pain). To ensure the behavioural task performance of the patients fulfilled this criterion, the ratings of each patient’s pain was evaluated based on a constructed parameter PR (see Supplementary Figure 1). See Supplementary Figures 2 and 3 for the detailed rating time courses of each session for each subject.

**Figure 1.**
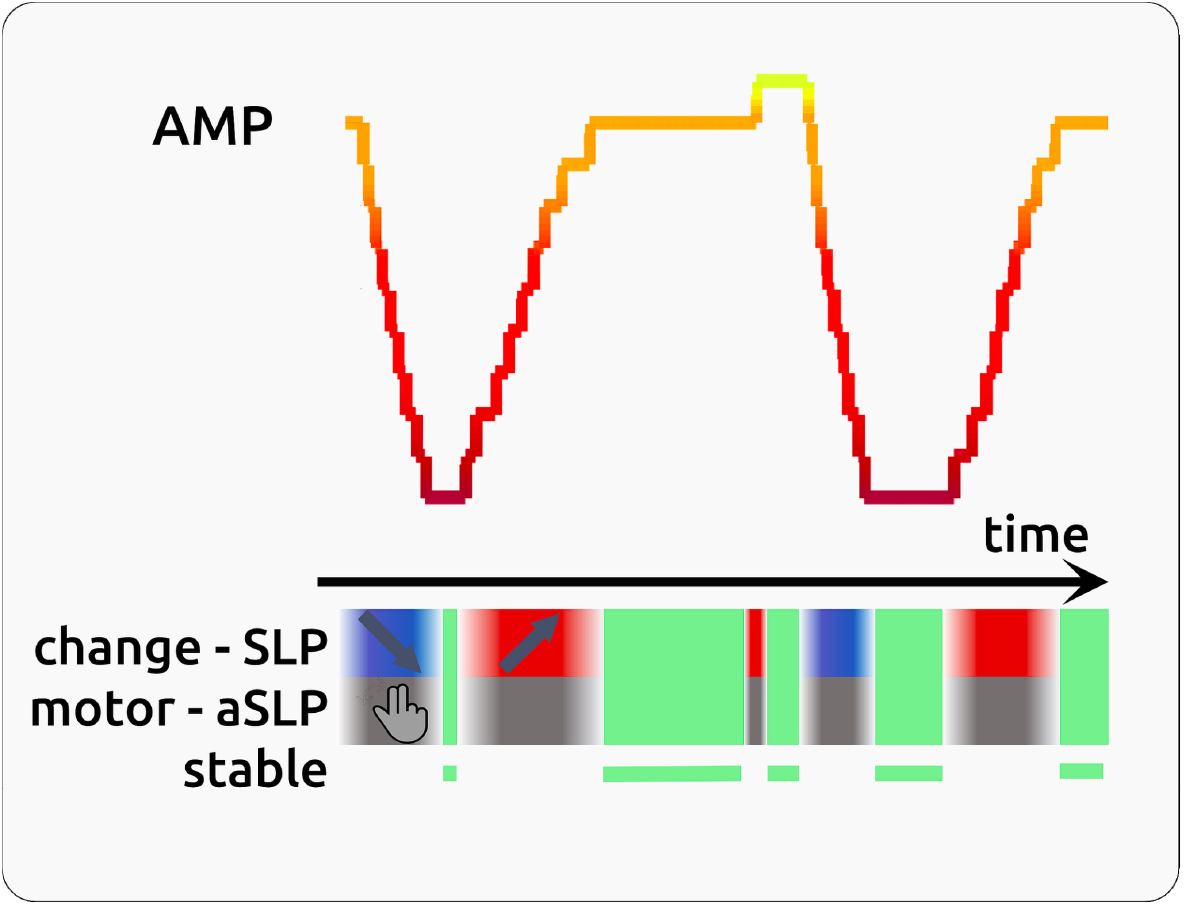
Schematic illustration of a hypothetical 3 min fluctuating time course of pain rating. The variable pain rating is colour-coded in red (low pain) to yellow (high pain). The balanced design ensures a similar amount of phases with rising pain (change - red) and falling pain (change - blue). Phases of stable pain are highlighted in light green. Slider movements (grey) are neither bound to pain amplitude (high, low) nor to change direction (rising, falling).

### Data processing - imaging data

Functional MRI data were preprocessed using FSL (Version 5.0.10) ^28^, which included removal of non-brain data (using brain extraction), slice time correction, head motion correction, B0 unwarping, spatial smoothing using a Gaussian kernel of FWHM (full width at half maximum) 6 mm, a nonlinear high-pass temporal filtering with a cutoff of 400 s, and spatial registration to the Montreal Neurological Institute (MNI) template. The data were further semi-automatically cleaned of artefacts with MELODIC ^29^. Artefact-related components were evaluated according to their spatial or temporal characteristics and were removed from the data following the recommendations in ^30,31^. The average number of artefact components for CM was 40±6 and for CBP 49±8. We deliberately did not include any correction for autocorrelation, neither for the processing of the imaging data nor for the processing of the pain rating time course as this step has the potential to destroy the natural evolution of the processes we aim to investigate (see PALM analysis below).

#### Regression and Scrubbing

Outliers in the fMRI data are labelled based on the definition of framewise displacement (FD) and DVARS following ^32^. A volume is defined as an outlier if it exceeds one of the following criteria: FD ≥ 0.2mm or DVARS ≥ (the 75th percentile + 1.5 times the interquartile range). Outliers are marked for each subject and then included in the regression as a vector of zeros (non-outlier) and ones (outlier). Each ‘window’ in the sliding window analysis (see below) was additionally scrubbed for outliers using Grubbs’ test ^33^; windows with more than two outliers are excluded from further analysis, including the preceding and following one. Windows with a correlation coefficient r >0.99 were omitted because these high correlations were mainly driven by outliers. The average percentage of scrubbed timepoints for all region pairs was 0.3±0.6 % for CBP and 0.3±0.4% for CM. MCFLIRT motion parameters, squared motion parameters, their temporal derivatives, and the squared temporal derivatives (total of 24 regressors) were included as motion regressors. The pain rating vector (AMP), the rate of change vector (SLP), and the absolute rate of change vector (aSLP) convolved with the hemodynamic response function were also included as regressors.

#### Cortical parcellation

We pursued a whole-brain parcellation approach ^34^ in order to assess every cortical connection that contributes to pain relief. Glasser’s parcellation subdivides the brain into 360 brain regions, 180 for each hemisphere of the brain; an additional 22 subcortical areas were added for a more detailed analysis (see Supplementary Table 3). To investigate the role of involvement of the cerebellum in pain processing, an additional 26 regions in the cerebellum (see Supplementary Table 4 for exact MNI coordinates) were added to the analysis.

#### Sliding-window connectivity measures

The time course of cortical activity for each of the 408 regions was computed using a principal component analysis (PCA). The first principal component was taken for further analysis. The ongoing connectivity between two brain regions was determined by Pearson’s correlation coefficient over sliding windows of 10 data points (t = TR*10= 15s). An offset ranging from -4 to 4 data points and a shift of one data point was used to account for a potential delay of information transfer between these regions. As a result we obtained a fluctuating time course covering 26 min of connectivity for each of the 408 seed regions with all other regions. The total number of analysed connections was 82824; each connection comprised 9 shifted time courses (-4 s to 4 s). Fisher’s Z-transformation was used for the normalisation of the correlation coefficients.

For the relationship between the correlation strength of two regions and the pain ratings, the rating vector was shifted between -15 s and 20 s in steps of 1 s (36 steps) against the connectivity time course. This procedure accounts for the unknown timing of cortical processing in reference to the rating and allows for some variability in the cortical response across brain regions; some ongoing cortical processes may influence later changes in pain ratings, other processes are directly related to the rating behaviour, or are influenced by the rating process and are occurring afterwards. The variable timing of the cortical processes are intermingled and we do not interpret the timing aspects any further. The same steps were applied for the created rate of change and absolute rate of change vector.

In order to assess cortical processes related to increasing and decreasing pain, the rate of change vector was calculated and the same shifting from -15 s to 20 s as for the amplitude time course was applied. Increasing pain is represented by a positive slope; decreasing pain by a negative slope. A vector of the absolute slope value represents a time course indicating the slider movement irrespective of the direction of the movement or the direction of the pain intensity change and was shifted as well.

Introducing these additional variables enabled us to disentangle the distinct connectivity patterns of pain intensity encoding (ongoing amplitude of pain rating - AMP) from brain processes related to the sensing of rising and falling pain (ongoing slope of pain rating - SLP) as well as from motor-related connectivity and decision making (ongoing absolute slope of pain ratings - aSLP).

### Statistical analysis - imaging data

Using Linear Mixed Effects models (LME; MixedModels.jl package in Julia) ^35^, we aimed to determine the relationship between fluctuating pain intensity and the fluctuating cortical connectivity separately for each pair of brain regions. The fluctuating connectivity of a particular pair is modelled through the time course of the three variables (AMP, SLP, aSLP) derived from the pain ratings (Figure 1).

In the description below, the statistical model is expressed in Wilkinson notation ^36^; the included fixed effects (connectivity ~ AMP + SLP + aSLP) describe the magnitudes of the population common intercept and the population common slopes for the relationship between cortical data and the intercept and these 3 variables. The added random effects (e.g. AMP - 1 | session) model the specific intercept differences for each recording session (e.g. session specific differences in pain levels or echo-planar image signal intensities):

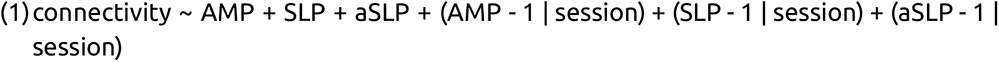

The common slope and the common intercept represent the group-wise fixed effects, which are the parameters of main interest in the LME. Hence, the model has 4 fixed effects parameters (common intercept, AMP, SLP, aSLP) and 234 (3*78) random effect parameters with related variance components of AMP random slopes, SLP random slope, aSLP random slope, and additive unexplained error. Each model was computed 36 times along the time shifts of the rating vector (-15 s to 20 s in steps of 1 s, see above). This procedure results in 36 t-values for each modality (AMP, SLP, and aSLP) and connection. For each modality the highest absolute t-values of the fixed effect parameters between -15 s and 20 s were extracted. The statistical model also included the information of the direction of change of pain intensity (SLP: encoded as -1 as shown in blue boxes for negative slope; encoded as 0 for stable phases of pain, and encoded as 1 as shown in red boxes for increasing pain). The task also required moving the potentiometer slider (aSLP: encoded as 1 as shown in grey for motor phases, and encoded as 0 for stable phases that did not require a motor process). Brain estimates of amplitude (AMP) should be independent irrespective of whether a data point originates from rising, stable or falling time points of the rating time course. In a similar vein, each slider movement involves a prior decision making and motor-related connectivity. These processes (SLP, aSLP) occur concomitant to the encoding of pain intensity (AMP) but are functionally, temporally and statistically independent.

### Correcting statistical testing - surrogate data

All statistical tests were corrected for multiple testing (connections, time shifts, rating shifts) and autocorrelation in the behavioural data: we created 1000 surrogate time courses using the IAAFT algorithm (Iterative Amplitude Adjusted Fourier Transform) from the original rating data, which were uncorrelated to the original rating data but had the same autocorrelation structure as the original data ^37^. Using surrogate data, the entire LME analysis was repeated 1000 times for the vectors (AMP, SLP, ASLP) with zero shift, resulting in 1000 whole-brain statistical maps for AMP, SLP and aSLP, respectively. From each map the highest absolute t-values of each repetition across the whole volume was extracted. This procedure resulted in a right-skewed distribution of 1000 values for each condition. Based on the distributions of 1000 values (for AMP, SLP, aSLP), the statistical thresholds were determined using the “palm_datapval.m” function publicly available in PALM ^38,39^.

### Analysis of individual and group connectivity maps

We investigated the individual connectivity confusion maps of endogenous pain encoding separately for each participant across all of their recordings. In order to assess whether the pattern of connectivity resembles the map of the group statistics, we correlated the connectivity of the group maps with the connectivity of the single-patient maps. The data were restricted to connections with an absolute value of t>0.75*PALM-threshold in the group statistics. We “normalised” the group map and single-patient maps with the following procedure: the pre-selected absolute t-values were ranked and equidistant numbers between 1 and 1000 were given to each included connections. Connections with negative t-values were given back their negative sign. Separately for each patient, the analysis was further restricted to connections for which the LME had converged. Spatial correlations using Kendall’s τ (tau) coefficients were computed for each patient’s connectivity map with the group connectivity map.

To visualise the connections between all significant region pairs, confusion matrices for all 408 regions were created as well as circle plots (NeuroMArVL, Brain Data Viewer, 2015). The weblinks to each plot are provided in the figure legends.

## Results

Here, we investigated the changes of cortical connectivity in relation to different aspects of the perception of ongoing chronic pain. We investigated whether correlated, anti-correlated, or uncorrelated connections reflect the perception of high pain states. Please note that we considered certain cases as uninterpretable. In these cases we would find changes of pain perception were relying on the entire range of connectivity, e.g. when low pain states were related to negatively correlated data, middle pain states to zero correlations, and high pain states to positive correlations. Consequently, we did not find any largely anti-correlated connectivity to be significantly related to pain (see schematic overview in Supplementary Figure 5). All findings were corrected for multiple testings (PALM, p<0.05).

### Connectivity pattern for the encoding of pain intensity across all CBP patients (AMP)

Across all subjects and sessions we found 2 connections to be positively related to the intensity of the endogenous pain (Figure 2); these pairs of brain regions, consisting of connections between cerebellar and parietal brain regions, showed a higher connectivity with higher levels of pain intensity.

**Figure 2.**
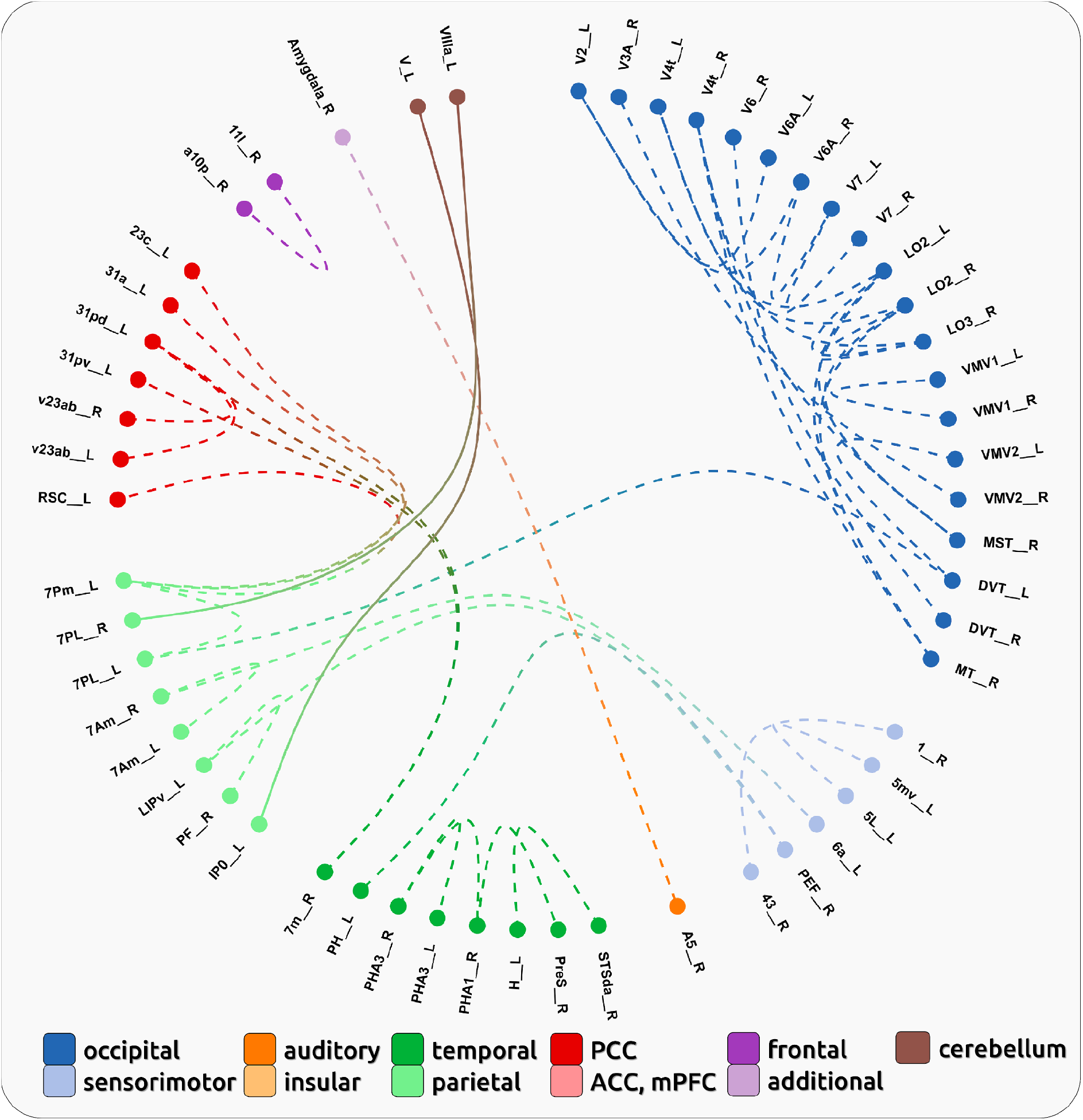
Connectivity pattern for the encoding of pain intensity across all CBP patients (AMP). Dashed lines indicate negative relationships with pain intensity; solid lines indicate positive relationships with pain intensity ([link](https://immersive.erc.monash.edu/neuromarvl/?save=44abd100-0b92-4af7-b76e-6b656d64c451_95.91.255.148)).

In addition we found 41 pairs of brain regions that exhibited a negative relationship between cortical connectivity and pain intensity (Figure 2). These pairs showed a lower connectivity with higher levels of experienced pain. One pattern of connectivity is related to the occipital cortex, where we found an overall disconnection within the lobe for higher pain states. Further disruptions of connectivity have been observed in limbic (hippocampal, parahippocampal), cingulate (BA23 and BA31), and somato-sensory areas, as well as in the precuneus (BA7).

Please note, that all 43 significant connections exhibited pain-related connectivity changes for positively correlated brain regions, i.e. the entire statistics largely relied on a sliding window of positive correlation coefficients. There was no significant effect for connections that relied on anti-correlated brain regions. Regions with more than one connection are listed in Supplementary Table 5. The confusion matrix for all connections is shown in Figure 4.

### Connectivity pattern for the encoding of the change of pain intensity across all CBP patients (SLP)

Across all subjects and sessions we found 9 connections to be positively related to the direction of change of the endogenous pain intensity (Figure 3). These pairs of brain regions showed increasing connectivity with rising pain intensity and include interhemispheric and left-lateralised intrahemispheric occipital connections, as well as connections between right angular and somatosensory regions.

**Figure 3.**
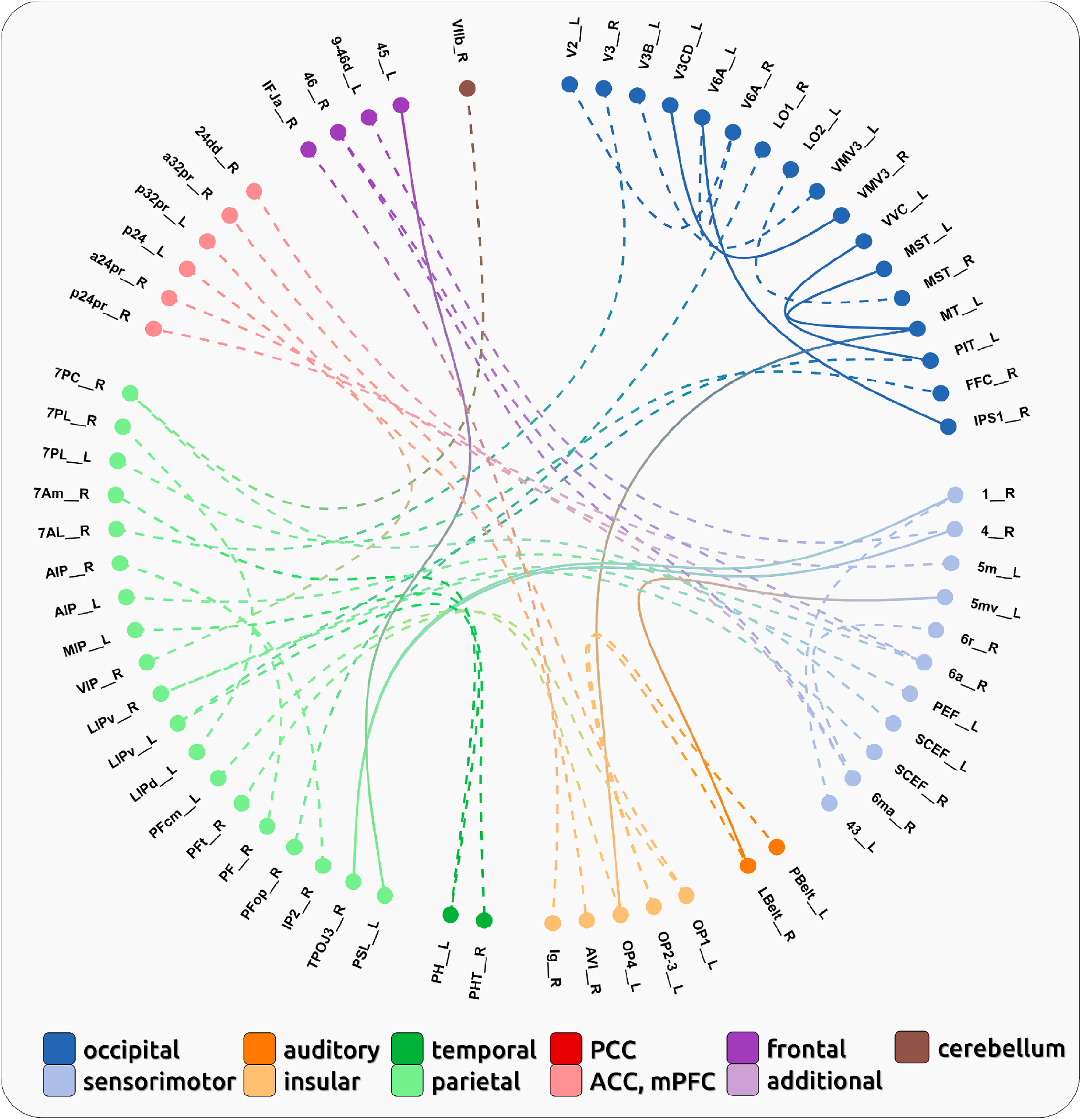
Connectivity pattern for the encoding of the change of pain intensity across all CBP patients (SLP). Dashed lines indicate negative relationships with rising pain intensity; solid lines indicate positive relationships with rising pain intensity ([link](https://immersive.erc.monash.edu/neuromarvl/?save=064d1a1b-17d8-4c55-8340-834b363d3906_95.91.255.148)).

**Figure 4.**
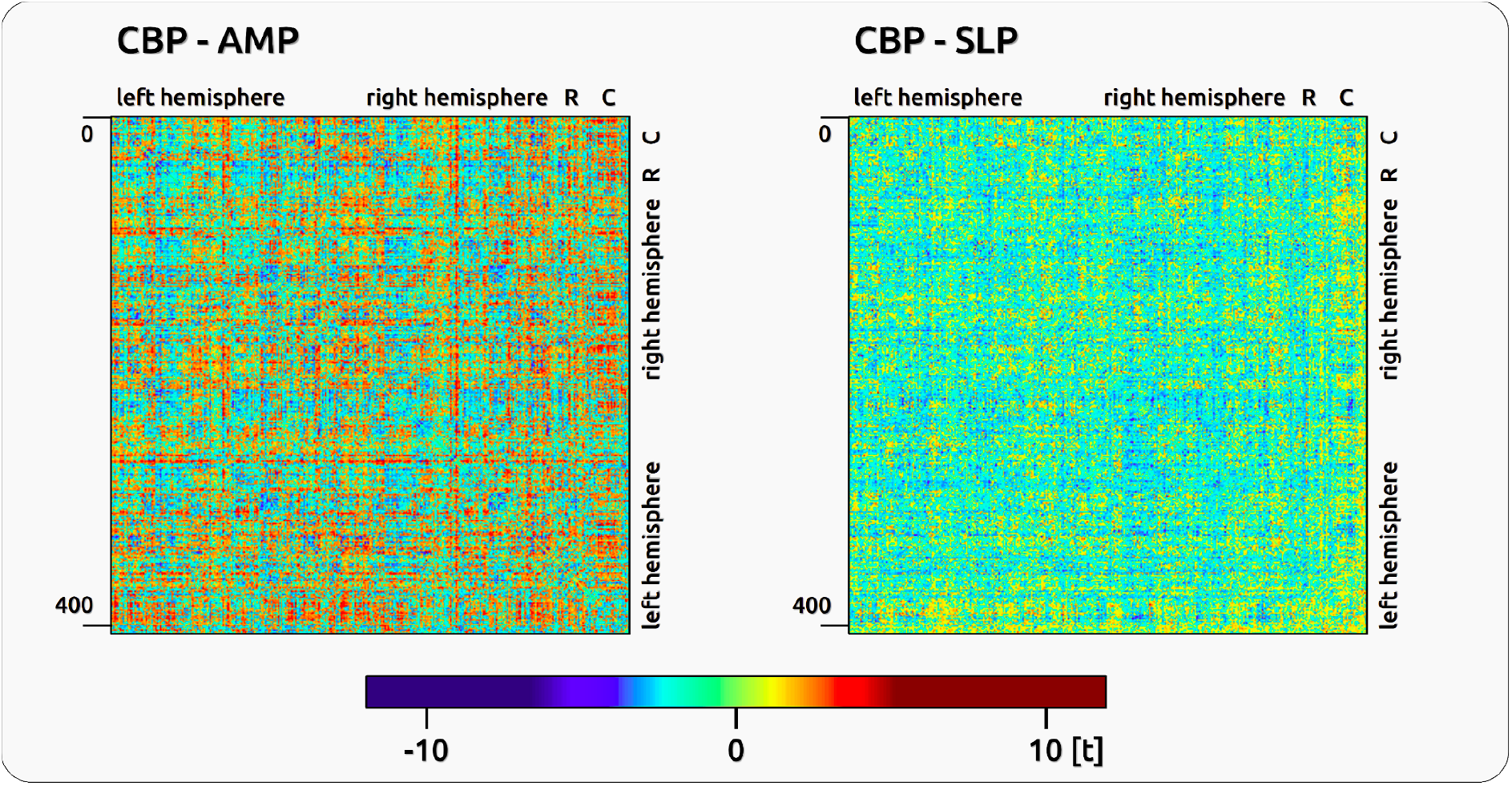
Confusion matrices for the encoding of AMP (left) and SLP (right) for all CBP subjects. Confusion matrices for AMP and SLP for all 408 region pairs given in t-values: 1-180: left hemisphere, 181-360: right hemisphere; 361-381: R: additional regions; 382-408: C: Cerebellum. AMP shows ~70% negative and ~30% positive significant t-values; the confusion matrix for SLP shows even more negative (~74%) than positive (~26%) ones.

The majority of the significantly connected brain regions exhibited a negative relationship between cortical connectivity and pain intensity (Figure 3). These 35 pairs showed decreasing connectivity with increasing levels of experienced pain. In other words, increasing pain is predominantly bound to a progressive loss of connectivity. This applies to connections that involve left parietal opercular regions, right insular regions, as well as large parts of the parietal, cingular and motor cortices.

Again, the entire statistics largely relied on a sliding window of positive correlation coefficients. There was no significant effect for connections that relied on anti-correlated brain regions. Regions with more than one connection are listed in Supplementary Table 6.

### Connectivity pattern for the encoding of pain intensity across all CM patients (AMP)

We found two connections that exhibited a significant effect for the encoding of pain intensity in CM. Decreasing connectivity between left motor regions (6d and 43) and increasing connectivity between the anterior insula and the orbitofrontal cortex indicate increasing intensity of CM (Figure 5). The confusion matrix for all connections is shown in Figure 7.

**Figure 5.**
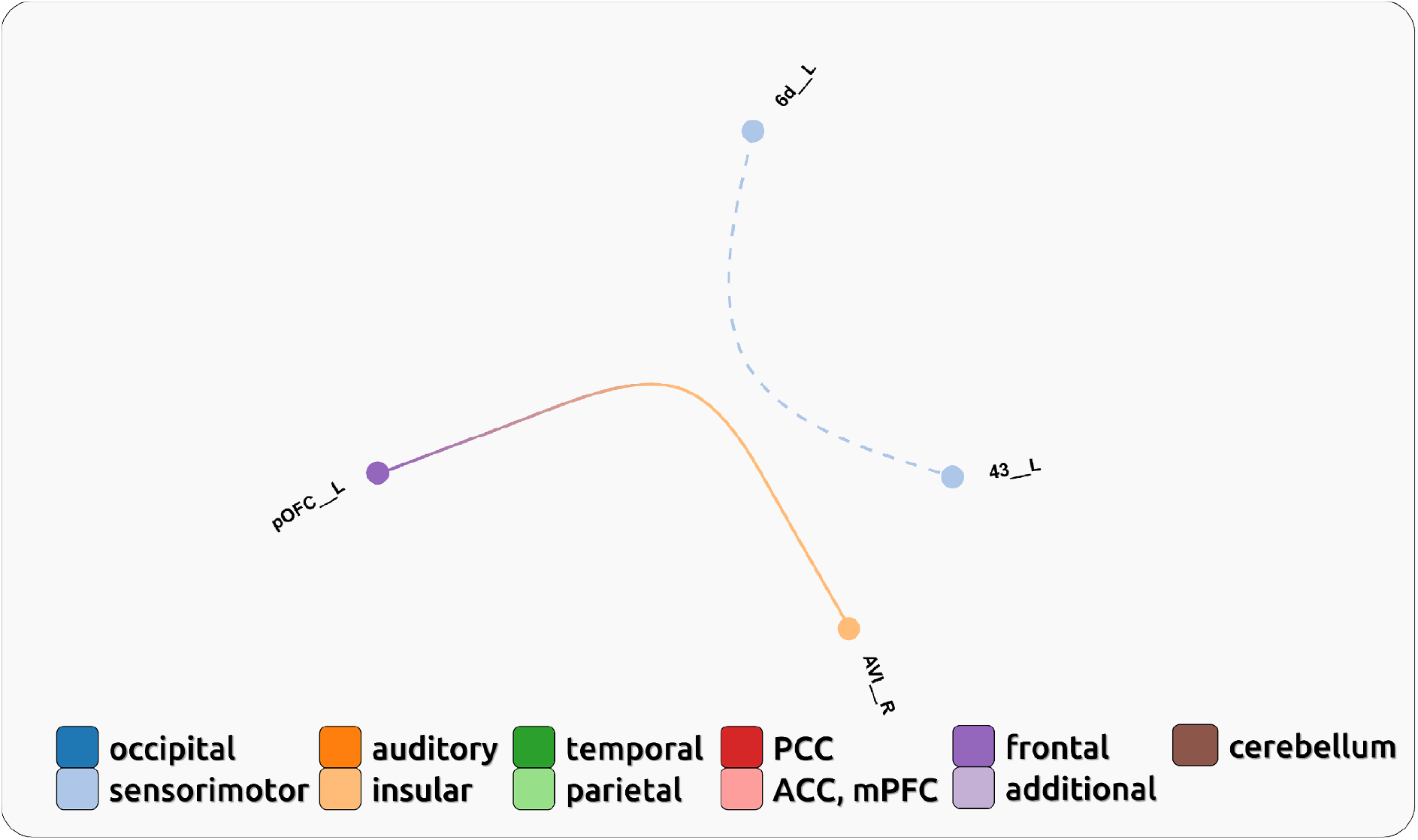
Connectivity pattern for the encoding of pain intensity changes across all CM patients (SLP). Dashed lines indicate negative relationships with rising pain intensity; solid lines indicate positive relationships with rising pain intensity ([link](https://immersive.erc.monash.edu/neuromarvl/?save=51f5ff51-c066-4c55-9758-6548fce2cdb9_95.91.255.148)).

### Connectivity pattern for the encoding of the change of pain intensity across all CM patients (SLP)

Across all subjects and sessions we found 4 connections to be positively related to the intensity of the endogenous pain (Figure 6); these pairs of brain regions, e.g. consisting of connections between temporal regions with the left insula and the hippocampus, showed increasing connectivity with higher levels of pain intensity.

**Figure 6.**
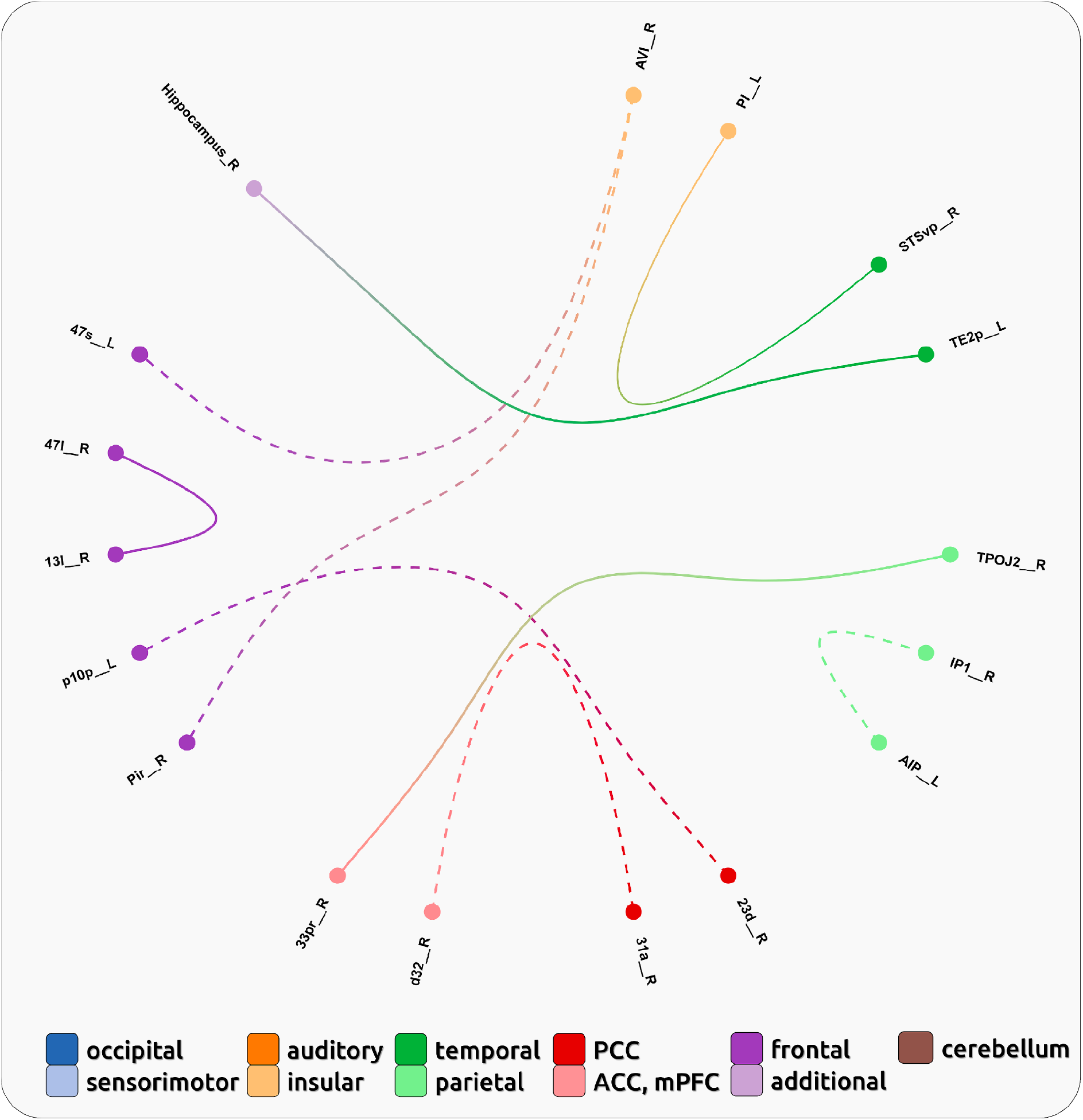
Connectivity pattern for the encoding of pain intensity across all CM patients (AMP). Dashed lines indicate negative relationships with rising pain intensity; solid lines indicate positive relationships with rising pain intensity ([link](https://immersive.erc.monash.edu/neuromarvl/?save=919a637c-5cef-4b34-aa18-fa718cfd728b_95.91.237.177)).

**Figure 7.**
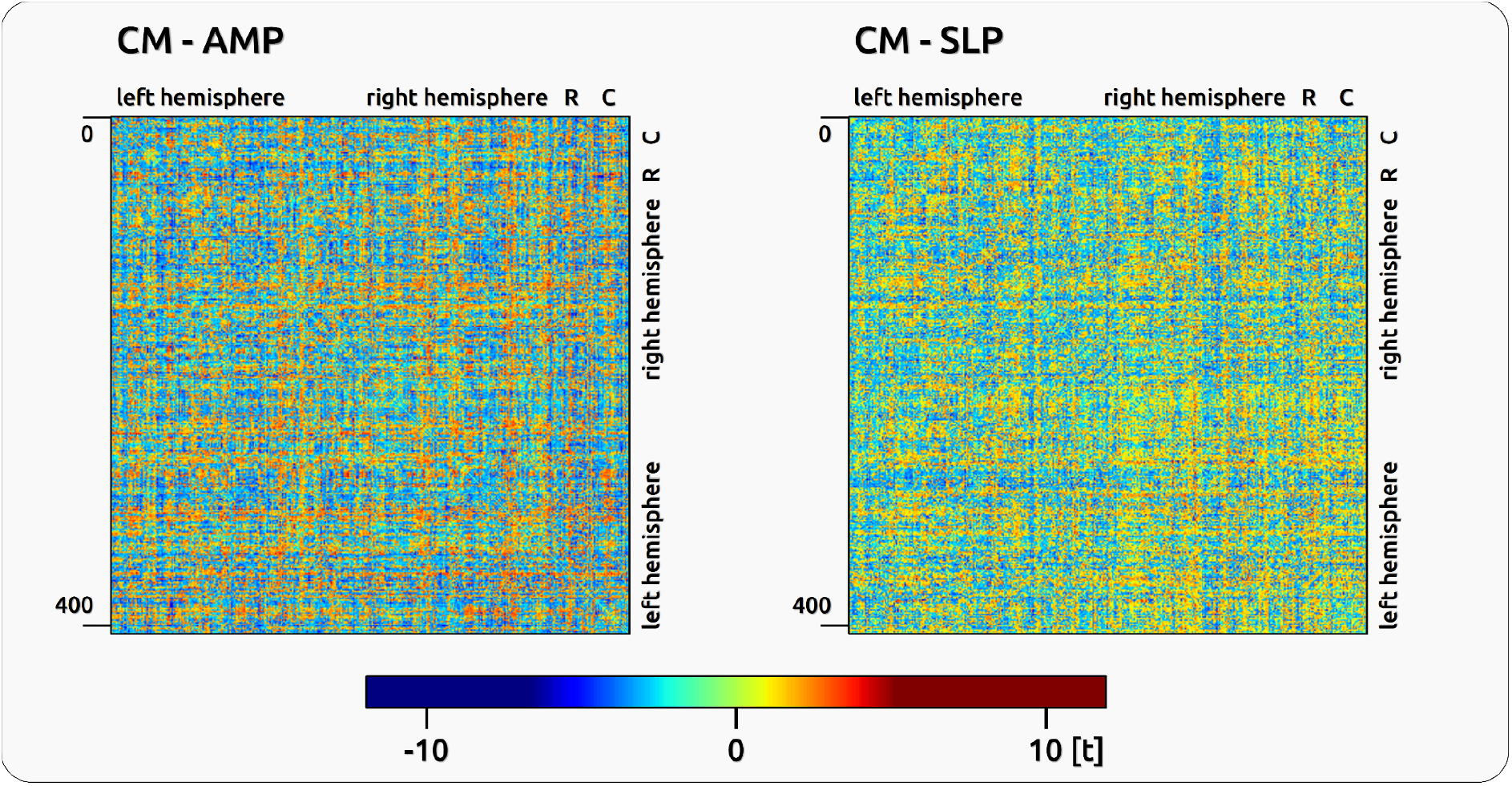
Confusion matrices for the encoding of AMP and SLP for all CM subjects. Confusion matrices for AMP and SLP for all 408 region pairs given in t-values: 1-180: left hemisphere, 181-360: right hemisphere; 361-381: additional regions (R); 382-408: Cerebellum (C). The confusion matrix for AMP shows 25% positive significant t-values, and 75% negative ones, compared to SLP which shows more positive (~58%) than negative (~42%) ones.

In addition we found 5 pairs of brain regions that exhibited a negative relationship between cortical connectivity and pain intensity (Figure 6). These pairs showed decreasing connectivity with lower levels of experienced pain, e.g. between the anterior insular cortex and orbitofrontal areas, as well as between the PCC and frontal and ACC regions.

### Individual patterns of pain encoding

The correlation between individual maps and the group results were calculated with Kendall’s tau and listed in Supplementary Table 7 for CBP and CM. The mean correlation between the individual AMP maps and the group AMP map for CBP/CM is tau = 0.09/0.07, tau = 0.03/0.05 for the SLP variable and tau = 0.09/0.1 for the aSLP variable. The number of all significant connections for CBP and CM for every subject is shown in Supplementary Table 8. Due to the large within-group variation and the finding that the group statistics does not represent a single individual, we refrained from computing any contrast between both experimental groups as the meaningfulness and interpretability of such a contrast would be very limited.

## Discussion

Here, we investigated the changes of cortical connectivity in relation to complementary aspects of the individual perception of ongoing chronic pain. We revealed positive and negative relationships between pain perception and positively correlated brain regions, but did not find any effect for anti-correlated (i.e. suppressing) brain regions. As a result, distinct patterns of pain-related connectivity for CBP and CM for the encoding of the magnitude of pain intensity as well as for the encoding of the change of direction of pain intensity (increasing vs. decreasing pain) were revealed. Overall, our results resemble recent findings of reduced cortical connectivity for higher pain states ^40^. The analysis of the repeated single-subject connectivity maps suggests a more complex picture, indicating a unique pattern of pain-related cortical processing for each patient. The group findings will be discussed in a traditional fashion but the single subject analysis suggests that none of the group findings apply to the cortical processing of a single subject. The findings suggest rather qualitative than gradual and quantitative differences between pain patients.

### Connectivity pattern encoding pain perception in CBP

#### Encoding of pain intensity

Overall, we found a predominantly negative relationship between cortical connectivity and high and rising pain in CBP. These findings reflect the severe impact of chronic pain on perception and cognitive functioning and are supportive of previous studies reporting how the experience of pain suppresses, inhibits, and impairs cortical processes (reviewed in ^41^.

Our findings on intra-occipital pain-intensity related disruptions are in line with the observations of suppressed occipital activity through applied pain ^42,43^ and the finding of disrupted visual network connectivity in chronic pain ^19^. Similar to our results, disrupted connectivity between the PCC and parietal regions was previously reported for acute tonic and chronic orofacial pain ^44^. Furthermore, impaired parahippocampal and hippocampal functioning as observed in chronic back pain has been discussed in the context of depression ^45,46^, biased memory ^47,48^, pain memory ^45,49^, and the transition from episodic to chronic pain ^50,51^.

Besides disrupted connectivity, we also found a positive relationship for cerebellar and parietal connectivity with increasing pain. The contribution of the cerebellum to the cortical processing of pain has been discussed in various contexts ^52^, including emotion, cognition, and motor functions, and has been suggested to integrate the multi-faceted aspects of the pain experience ^53^. Consequently, suffering from long-lasting pain may have caused structural changes in the cerebellum. For CBP, alterations of the cerebellar grey matter density were observed compared to healthy subjects ^54,55^. Both the cerebellum and the connected parietal regions have been associated with a top-down attentional direction of pain intensity features ^56^.

#### Encoding of the direction of pain intensity changes

We contrasted periods of rising pain with periods of falling pain. This contrasts controls for motor-related connectivity, the current level of pain, and decision making processes, as these processes equally occur during rising and falling pain. As a result, we observed a wide-spread disruption of cortical circuits with rising pain; the connectivity was significantly higher during falling pain for a number of cortical connections. This applied to the connectivity between brain regions commonly associated with pain perception, i.e. insular cortex to perigenual ACC ^57^. We have previously reported lower connectivity in higher pain states for regions that are known to be involved in the encoding of pain ^40^.

In addition, we also observed a disruption within the parietal and the cingulate cortices. The parietal disruption may reflect the deviant processing within the default mode network, which has been previously reported ^11,44,58^.

### Connectivity pattern encoding pain perception in CM

#### Encoding of pain intensity

For CM, the group statistics showed only 2 connections that represent the encoding of pain intensity. Among these connections we found an increasing connectivity with higher pain intensities between the right anterior insular and the left orbitofrontal cortex. The orbitofrontal cortex ^59,60^ and anterior insular cortex ^42^ have been consistently found to be involved in pain processing. The comparably low number of connections may represent the complexity of the cortical processing in chronic migraine ^61,62^. Indeed, most patients that were included in the present study exhibited unique patterns of pain-related cortical connectivity that do not match the overall pattern of the group statistics (see below).

#### Encoding of the direction of pain intensity changes

There are a number of connections that represent the rising and falling pain in CM. We found an involvement of the right anterior insula that is connected to orbitofrontal areas; one of these frontal regions has been previously related to migraine attacks ^63^ and placebo modulation (Figure 2C in ^64^). The pain-modulated connectivity between the anterior and the posterior cingulate cortex may reflect the activity of the DMN ^11,44,65^.

Superior temporal regions, which we found to be positively connected to the insula, have been shown to be affected in episodic migraine ^66^ and pain memory ^67^. This may also apply to the connection between the hippocampus and an inferior temporal area. Again, the majority of the regions exhibit a disrupted connectivity with rising pain.

Our findings, however, are not directly comparable to previous studies as we have assessed the non-stationary aspects of the cortical network. Here, we investigated the pain-dependent within-subject dynamic fluctuations irrespective of the general amplitude of the BOLD oscillations. This perspective on the data considers and contrasts only the rising and falling periods.

### Individual patterns of pain intensity encoding in single patients

The cornerstone of the present study is assessing each patients’ pain processing profile. To examine these unique patterns of pain-related connectivity, each patient underwent repeated recordings in order to create sufficient and reliable data. Individual connectivity maps were generated for each patient, all of which varied significantly from the connectivity pattern of the group. Indeed as a large number of connections were variable, we decided against a direct comparison between patient groups.

To regard individual variations within sensory data merely as noise may in fact hinder our understanding of sensory processing. Such noisy aspects of data may reflect pain processing, the experience or pain, or even the individual coping strategies of the patient, all of which may influence the success of chronic pain treatments. Likewise, the importance of examining individual patterns of brain connectivity and structure has previously been promoted ^6^other studies which address individual differences and single patient effects have also reported variability ^68–70^. Although group statistics might suggest otherwise, individual parameters such as gamma oscillations in tonic and chronic pain reveal that pain is not encoded by gamma activity in all study participants ^71,72^. Such immense variability between individual pain signatures instead indicates qualitative differences between pain patients rather than quantitative differences ^73^. Variations in cortical processing are veritably in line with the clinical picture of pain diseases; individual patients each express a composite of characteristics unique to them ^74^. Yet, one must not exclude the possibility that variables such as pain duration and intensity, current medication, or indeed psychological parameters and subtypes of pain disease may modulate a number of aspects for a specific individual. For the present study, however, we have interpreted qualitatively variable patterns being modulated by such variables as improbable and thus excluded comparisons between behavioural and cortical data. By assessing individual pain signatures, we may facilitate more accurate assessment of chronic pain conditions, which is in line with recent developments to enhance and promote individually-tailored treatments in medicine ^75,76^.

## Conclusion

The experimental setup aimed to reflect the dynamically evolving cortical connectivities related to the subjective experience of pain. However, none of the single subject connectivity patterns, assessed in repeated sessions, resembled the patterns obtained by group statistics. The results suggest that individual chronic pain patients exhibit qualitatively distinct signatures of cortical connectivity; this applies to chronic back pain as well as to chronic migraine.

Consequently, the present findings argue against a common biomarker for the subjective experience of chronic pain that is based on dynamic connections. This is in line with the experience of clinicians; each patient can be characterised by a unique personality, various combinations of symptoms, and a broad range of treatment success. The findings support recent developments for a more personalised medicine. Our data would support an individually-tailored therapeutic approach in clinical settings.

Our study can open a new window for the study of processes subserved in the human brain: research studies need to show reliable observations for both group statistics and individual patterns. This expands the current view on the replication crisis in neuroscience.

## Acknowledgements

This work was supported by the Deutsche Forschungsgemeinschaft (SCHU2879/4-1). We thank Dr Vrginia Flanagin for her comments on the methods and Dr Stephanie Irving for copy-editing the manuscript.

## Data availability

The data that support the findings of this study are available from the corresponding author upon reasonable request.

