## Supplementary Figure 1 for "Individually Unique Dynamics of Cortical Connectivity Reflect the Ongoing Intensity of Chronic Pain"

| | |
|---|---|
| Journal: | *Brain* |
| Manuscript ID | BRAIN-2021-01280 |
| Manuscript Type: | Original Article |
| Date Submitted by the Author: | 29-Jun-2021 |
| Complete List of Authors: | Mayr, Astrid; Ludwig Maximilians University Munich, Neurology<br>Jahn, Pauline; Klinikum der Universität München, Klinik und Poliklinik für Radiologie<br>Deak, Bettina; Ludwig Maximilians University Munich, Neurology<br>Stankewitz, Anne; Klinikum der Universität München, Neurologische Klinik und Poliklinik<br>Devulapally, Vasudev; Ludwig Maximilians University Munich, Neurology<br>Witkovsky, Viktor; Institute of Measurement Science SAS, Department of Theoretical Methods<br>Dietrich, Olaf; Ludwig-Maximilian University, Department of Clinical Radiology<br>Schulz, Enrico; Ludwig Maximilians University Munich, Department of Neurology |
| Methodology: | IMAGING |
| Subject area: | HEADACHE, PAIN |

SCHOLARONE™
Manuscripts

### Individually Unique Dynamics of Cortical Connectivity Reflect the Ongoing Intensity of Chronic Pain

Astrid Mayr[1,2], Pauline Jahn[2], Bettina Deak[2], Anne Stankewitz[2], Vasudev Devulapally[2], Viktor Witkovsky[3], Olaf Dietrich[1], Enrico Schulz[2,4]

[1] Department of Radiology, University Hospital, LMU Munich, 81377 Munich, Germany
[2] Department of Neurology, Ludwig-Maximilians-University Hospital Munich, 81377 Munich, Germany
[3] Department of Theoretical Methods, Institute of Measurement Science, Slovak Academy of Sciences, 841 04 Bratislava, Slovak Republic
[4] Department of Medical Psychology, Ludwig-Maximilians-University Hospital Munich, 81377 Munich, Germany

For Peer Review

**Corresponding author:**
Astrid Mayr
Ludwig-Maximilians-Universität München
Neurologische Klinik und Poliklinik
A: Fraunhoferstr. 20, 82152 Martinsried, Germany
O: +49 89 4400 74826
E:

ScholarOne, 375 Greenbrier Drive, Charlottesville, VA, 22901 Support (434) 964 4100

### Abstract

*Background.* Chronic pain diseases are characterised by an ongoing and fluctuating endogenous pain, yet it remains to be elucidated how this is reflected by the dynamics of ongoing functional cortical connections. The present study addresses this disparity by taking the individual perspective of pain patients into account, which is the varying intensity of endogenous pain.

For Peer Review

ScholarOne, 375 Greenbrier Drive, Charlottesville, VA, 22901 Support (434) 964 4100

### Introduction

Functional connectivity (FC) in the brain is a measure of spatio-temporal correlations of fluctuating time-series of distinct regions. It has become essential for exploring the functional network structure of both healthy and diseased brains [1–4] as alterations in functional connectivity patterns due to neurological or psychiatric disorders, as well as changes due to long-lasting chronic pain conditions, have been reported [5–7]. Functional connectivity in chronic pain conditions have been studied by analysing intrinsic cortical networks as well as by computing seed-based connections of one or more predefined brain regions.

For Peer Review

ScholarOne, 375 Greenbrier Drive, Charlottesville, VA, 22901 Support (434) 964 4100

visual network and S1 was reported, and connectivity strength was further negatively correlated with the disease duration.

There are a number of challenges that hamper the formation of a unified framework for the understanding of the cortical underpinnings of chronic pain disease and may explain the often contradicting findings. *First,* the cause of the common low-frequency fluctuation of BOLD activity, which gives rise to distinct intrinsic networks, is not yet entirely understood. The BOLD fluctuations, consisting of peaks and troughs, may be related to the continuously-changing pain intensity. This relationship has not been investigated so far and network fluctuations that are unrelated to the ongoing experience of pain would be difficult to interpret. *Second*, many studies in the neuroimaging literature lack reliable cortical data with repeated measurements for reproducible and longitudinal observations [6,20], which may explain the heterogeneity of the DMN findings. Therefore, extended and repeated recordings are required to disentangle a subject's stable signature from random momentary fluctuations of brain connectivity.

Here, we aimed to capture the individual endogenous pain experience for each subject. We utilised a whole-brain functional connectivity approach to identify coherent brain connectivity patterns of chronic pain encoding (CBP and CM). To this end, pain ratings were related to simultaneously obtained cortical data to characterise the fluctuating neuronal connectivity patterns that reflect the experience of chronic pain for each individual across several recordings. Multiple recordings for each individual enabled us to explore how well the results of group statistics reflect the individual processing of chronic pain.

For Peer Review

ScholarOne, 375 Greenbrier Drive, Charlottesville, VA, 22901 Support (434) 964 4100

### Material and Methods

#### Participants

The study included 20 patients diagnosed with chronic back pain (CBP - 16 female; aged 44±13 years) and 20 patients with chronic migraine (CM - 18 female; aged 34±13 years). All participants gave written informed consent. The study was approved by the Ethics Committee of the Medical Department of the Ludwig-Maximilians-Universität München and conducted in conformity with the Declaration of Helsinki.

For Peer Review

ScholarOne, 375 Greenbrier Drive, Charlottesville, VA, 22901 Support (434) 964 4100

functional MRI session. The screen was visible through a mirror mounted on top of the MRI head coil. Patients were asked to look only at the screen and to focus on their pain. The intensity and the changes of perceived pain had to be indicated as quickly and accurately as possible. To minimise head movement, foams were placed around the head and patients were told to lie as still as possible.

ScholarOne, 375 Greenbrier Drive, Charlottesville, VA, 22901 Support (434) 964 4100
For Peer Review

level, in order to mitigate potential effects of order (e.g. in case of continuously rising pain). To ensure the behavioural task performance of the patients fulfilled this criterion, the ratings of each patient's pain was evaluated based on a constructed parameter PR (see Supplementary Figure 1). See Supplementary Figures 2 and 3 for the detailed rating time courses of each session for each subject.

ScholarOne, 375 Greenbrier Drive, Charlottesville, VA, 22901 Support (434) 964 4100

##### Sliding-window connectivity measures

The time course of cortical activity for each of the 408 regions was computed using a principal component analysis (PCA). The first principal component was taken for further analysis. The ongoing connectivity between two brain regions was determined by Pearson's correlation coefficient over sliding windows of 10 data points (t = TR*10= 15s). An offset ranging from -4 to 4 data points and a shift of one data point was used to account for a potential delay of information transfer between these regions. As a result we obtained a fluctuating time course covering 26 min of connectivity for each of the 408 seed regions with all other regions. The total number of analysed connections was 82824; each connection comprised 9 shifted time courses (-4 s to 4 s). Fisher's Z-transformation was used for the normalisation of the correlation coefficients.

ScholarOne, 375 Greenbrier Drive, Charlottesville, VA, 22901 Support (434) 964 4100

(e.g. AMP - 1 | session) model the specific intercept differences for each recording session (e.g. session specific differences in pain levels or echo-planar image signal intensities):

ScholarOne, 375 Greenbrier Drive, Charlottesville, VA, 22901 Support (434) 964 4100

values were given back their negative sign. Separately for each patient, the analysis was further restricted to connections for which the LME had converged. Spatial correlations using Kendall's τ (tau) coefficients were computed for each patient's connectivity map with the group connectivity map.

ScholarOne, 375 Greenbrier Drive, Charlottesville, VA, 22901 Support (434) 964 4100

left-lateralised intrahemispheric occipital connections, as well as connections between right angular and somatosensory regions.

#### Discussion

Here, we investigated the changes of cortical connectivity in relation to complementary aspects of the individual perception of ongoing chronic pain. We revealed positive and

ScholarOne, 375 Greenbrier Drive, Charlottesville, VA, 22901 Support (434) 964 4100
For Peer Review

negative relationships between pain perception and positively correlated brain regions, but did not find any effect for anti-correlated (i.e. suppressing) brain regions.

ScholarOne, 375 Greenbrier Drive, Charlottesville, VA, 22901 Support (434) 964 4100

In addition, we also observed a disruption within the parietal and the cingulate cortices. The parietal disruption may reflect the deviant processing within the default mode network, which has been previously reported [11,44,58].

For Peer Review

ScholarOne, 375 Greenbrier Drive, Charlottesville, VA, 22901 Support (434) 964 4100

have also reported variability [68–70]. Although group statistics might suggest otherwise, individual parameters such as gamma oscillations in tonic and chronic pain reveal that pain is not encoded by gamma activity in all study participants [71,72]. Such immense variability between individual pain signatures instead indicates qualitative differences between pain patients rather than quantitative differences [73]. Variations in cortical processing are veritably in line with the clinical picture of pain diseases; individual patients each express a composite of characteristics unique to them [74]. Yet, one must not exclude the possibility that variables such as pain duration and intensity, current medication, or indeed psychological parameters and subtypes of pain disease may modulate a number of aspects for a specific individual. For the present study, however, we have interpreted qualitatively variable patterns being modulated by such variables as improbable and thus excluded comparisons between behavioural and cortical data. By assessing individual pain signatures, we may facilitate more accurate assessment of chronic pain conditions, which is in line with recent developments to enhance and promote individually-tailored treatments in medicine [75,76].

For Peer Review

**Conclusion**

The experimental setup aimed to reflect the dynamically evolving cortical connectivities related to the subjective experience of pain. However, none of the single subject connectivity patterns, assessed in repeated sessions, resembled the patterns obtained by group statistics. The results suggest that individual chronic pain patients exhibit qualitatively distinct signatures of cortical connectivity; this applies to chronic back pain as well as to chronic migraine.

Consequently, the present findings argue against a common biomarker for the subjective experience of chronic pain that is based on dynamic connections. This is in line with the experience of clinicians; each patient can be characterised by a unique personality, various combinations of symptoms, and a broad range of treatment success. The findings support recent developments for a more personalised medicine. Our data would support an individually-tailored therapeutic approach in clinical settings.

Our study can open a new window for the study of processes subserved in the human brain: research studies need to show reliable observations for both group statistics and individual patterns. This expands the current view on the replication crisis in neuroscience.

### Acknowledgements

This work was supported by the Deutsche Forschungsgemeinschaft (SCHU2879/4-1). We thank Dr Vrginia Flanagin for her comments on the methods and Dr Stephanie Irving for copy-editing the manuscript.

### Data availability

The data that support the findings of this study are available from the corresponding author upon reasonable request.

### Competing interests

The authors report no competing interests.

ScholarOne, 375 Greenbrier Drive, Charlottesville, VA, 22901 Support (434) 964 4100

For Peer Review

*Behaviour Research and Therapy*. 1995;33(3):335-343. doi:10.1016/0005-7967(94)00075-u
26. Schulz E, Stankewitz A, Witkovský V, Winkler AM, Tracey I. Strategy-dependent modulation of cortical pain circuits for the attenuation of pain. *Cortex*. 2019;113:255-266.
27. Penny WD, Friston KJ, Ashburner JT, Kiebel SJ, Nichols TE. *Statistical Parametric Mapping: The Analysis of Functional Brain Images*. Elsevier; 2011.
28. Jenkinson M, Beckmann CF, Behrens TEJ, Woolrich MW, Smith SM. FSL. *Neuroimage*. 2012;62(2):782-790.
29. Salimi-Khorshidi G, Douaud G, Beckmann CF, Glasser MF, Griffanti L, Smith SM. Automatic denoising of functional MRI data: combining independent component analysis and hierarchical fusion of classifiers. *Neuroimage*. 2014;90:449-468.
30. Kelly RE, Alexopoulos GS, Wang Z, et al. Visual inspection of independent components: defining a procedure for artifact removal from fMRI data. *J Neurosci Methods*. 2010;189(2):233-245.
31. Griffanti L, Salimi-Khorshidi G, Beckmann CF, et al. ICA-based artefact removal and accelerated fMRI acquisition for improved resting state network imaging. *Neuroimage*. 2014;95:232-247.
32. Power JD, Barnes KA, Snyder AZ, Schlaggar BL, Petersen SE. Spurious but systematic correlations in functional connectivity MRI networks arise from subject motion. *Neuroimage*. 2012;59(3):2142-2154.
33. Grubbs FE. Sample Criteria for Testing Outlying Observations. *Ann Math Stat*. 1950;21(1):27-58.
34. Glasser MF, Coalson TS, Robinson EC, et al. A multi-modal parcellation of human cerebral cortex. *Nature*. 2016;536(7615):171-178.
35. Bezanson J, Edelman A, Karpinski S, Shah VB. Julia: A Fresh Approach to Numerical Computing. *SIAM Rev*. 2015;59(1):65-98.
36. Wilkinson GN, Rogers CE. Symbolic Description of Factorial Models for Analysis of Variance. *Applied Statistics*. 1973;22(3):392. doi:10.2307/2346786
37. Schreiber T, Schmitz A. Improved Surrogate Data for Nonlinearity Tests. *Phys Rev Lett*. 1996;77(4):635-638.
38. Winkler AM, Ridgway GR, Webster MA, Smith SM, Nichols TE. Permutation inference for the general linear model. *Neuroimage*. 2014;92:381-397.
39. Winkler AM, Webster MA, Brooks JC, Tracey I, Smith SM, Nichols TE. Non-parametric combination and related permutation tests for neuroimaging. *Hum Brain Mapp*. 2016;37(4):1486-1511.
40. Schulz E, Stankewitz A, Winkler AM, Irving S, Witkovský V, Tracey I. Ultra-high-field imaging reveals increased whole brain connectivity underpins cognitive strategies that attenuate pain. *Elife*. 2020;9. doi:10.7554/eLife.55028
41. Bushnell MC, Ceko M, Low LA. Cognitive and emotional control of pain and its disruption in chronic pain. *Nat Rev Neurosci*. 2013;14(7):502-511.
42. Wager TD, Atlas LY, Lindquist MA, Roy M, Woo C-W, Kross E. An fMRI-based neurologic signature of physical pain. *N Engl J Med*. 2013;368(15):1388-1397.
43. Ploner M, Gross J, Timmermann L, Pollok B, Schnitzler A. Pain suppresses spontaneous brain rhythms. *Cereb Cortex*. 2006;16(4):537-540.
44. Alshelh Z, Marciszewski KK, Akhter R, et al. Disruption of default mode network dynamics in acute and chronic pain states. *Neuroimage Clin*. 2018;17:222-231.
45. Mokhtari T, Tu Y, Hu L. Involvement of the hippocampus in chronic pain and depression. *Brain Science Advances*. 2019;5(4):288-298.
46. Fasick V, Spengler RN, Samankan S, Nader ND, Ignatowski TA. The hippocampus and TNF: Common links between chronic pain and depression. *Neurosci Biobehav Rev*. 2015;53:139-159.
47. Berger SE, Vachon-Presseau É, Abdullah TB, Baria AT, Schnitzer TJ, Apkarian AV. Hippocampal morphology mediates biased memories of chronic pain. *Neuroimage*. 2018;166:86-98.
48. Perry C, Heathcote L, Thomas K, Borsook D, Simons L, Timmers I. The role of hippocampus and amygdala morphology in memory bias of learned fear in youth with chronic pain. *J Pain*. 2021;22(5):604.
49. Sandkühler J, Lee J. How to erase memory traces of pain and fear. *Trends Neurosci*. 2013;36(6):343-352.
50. Mansour A, Farmer M, Baliki M, Apkarian A. Chronic pain: The role of learning and brain plasticity. *Restor Neurol Neurosci*. 2013;32. doi:10.3233/RNN-139003
51. Mutso A, Radzicki D, Baliki MN, et al. Abnormalities in hippocampal functioning with persistent pain. *J Neurosci*. 2012;32(17):5747-5756.
52. Claassen J, Koenen LR, Ernst TM, et al. Cerebellum is more concerned about visceral than somatic pain. *J Neurol Neurosurg Psychiatry*. 2020;91(2):218-219.
53. Moulton EA, Schmahmann JD, Becerra L, Borsook D. The cerebellum and pain: passive

For Peer Review

integrator or active participator? *Brain Res Rev*. 2010;65(1):14-27.
54. Luchtmann M, Steinecke Y, Baecke S, et al. Structural brain alterations in patients with lumbar disc herniation: a preliminary study. *PLoS One*. 2014;9(3):e90816.
55. Ung H, Brown JE, Johnson KA, Younger J, Hush J, Mackey S. Multivariate classification of structural MRI data detects chronic low back pain. *Cereb Cortex*. 2014;24(4):1037-1044.
56. Lobanov OV, Quevedo AS, Hadsel MS, Kraft RA, Coghill RC. Frontoparietal mechanisms supporting attention to location and intensity of painful stimuli. *Pain*. 2013;154(9):1758-1768.
57. Tracey I, Mantyh PW. The cerebral signature for pain perception and its modulation. *Neuron*. 2007;55(3):377-391.
58. Loggia ML, Kim J, Gollub RL, et al. Default mode network connectivity encodes clinical pain: An arterial spin labeling study. *Pain*. 2013;154(1):24-33. doi:10.1016/j.pain.2012.07.029
59. Winston JS, Vlaev I, Seymour B, Chater N, Dolan RJ. Relative valuation of pain in human orbitofrontal cortex. *J Neurosci*. 2014;34(44):14526-14535.
60. Atlas LY, Bolger N, Lindquist MA, Wager TD. Brain mediators of predictive cue effects on perceived pain. *J Neurosci*. 2010;30(39):12964-12977.
61. Burstein R, Noseda R, Borsook D. Migraine: multiple processes, complex pathophysiology. *J Neurosci*. 2015;35(17):6619-6629.
62. Dodick DW. A phase-by-phase review of migraine pathophysiology. *Headache: the journal of head and face pain*. 2018;58:4-16.
63. Denuelle M, Fabre N, Payoux P, Chollet F, Geraud G. Hypothalamic activation in spontaneous migraine attacks. *Headache*. 2007;47(10):1418-1426.
64. Wager TD, Rilling JK, Smith EE, et al. Placebo-induced changes in FMRI in the anticipation and experience of pain. *Science*. 2004;303(5661):1162-1167.
65. Coppola G, Di Renzo A, Tinelli E, et al. Resting state connectivity between default mode network and insula encodes acute migraine headache. *Cephalalgia*. 2018;38(5):846-854.
66. Schwedt TJ, Chong CD, Chiang C-C, Baxter L, Schlaggar BL, Dodick DW. Enhanced pain-induced activity of pain-processing regions in a case-control study of episodic migraine. *Cephalalgia*. 2014;34(12):947-958.
67. Houde F, Martel M, Coulombe-Lévêque A, et al. Perturbing the activity of the superior temporal gyrus during pain encoding prevents the exaggeration of pain memories: A virtual lesion study using single-pulse transcranial magnetic stimulation. *Neurobiol Learn Mem*. 2020;169:107174.
68. Gordon EM, Laumann TO, Gilmore AW, et al. Precision Functional Mapping of Individual Human Brains. *Neuron*. 2017;95(4):791-807.e7.
69. Greene DJ, Marek S, Gordon EM, et al. Integrative and Network-Specific Connectivity of the Basal Ganglia and Thalamus Defined in Individuals. *Neuron*. Published online 2019. doi:10.1016/j.neuron.2019.11.012
70. Marek S, Siegel JS, Gordon EM, et al. Spatial and Temporal Organization of the Individual Human Cerebellum. *Neuron*. 2018;100(4):977-993.e7.
71. Schulz E, May ES, Postorino M, et al. Prefrontal Gamma Oscillations Encode Tonic Pain in Humans. *Cereb Cortex*. 2015;25(11):4407-4414.
72. May ES, Nickel MM, Dinh ST, et al. Prefrontal gamma oscillations reflect ongoing pain intensity in chronic back pain patients. *Hum Brain Mapp*. 2019;40(1):293-305.
73. Zadelaar JN, Weeda WD, Waldorp LJ, Van Duijvenvoorde ACK, Blankenstein NE, Huizenga HM. Are individual differences quantitative or qualitative? An integrated behavioral and fMRI MIMIC approach. *Neuroimage*. 2019;202:116058.
74. Smith HS. *Current Therapy in Pain*. Elsevier Health Sciences; 2009.
75. Mun CJ, Suk HW, Davis MC, et al. Investigating intraindividual pain variability: methods, applications, issues, and directions. *Pain*. 2019;160(11):2415-2429.
76. Ott PA, Hu Z, Keskin DB, et al. An immunogenic personal neoantigen vaccine for patients with melanoma. *Nature*. 2017;547(7662):217-221.

For Peer Review

ScholarOne, 375 Greenbrier Drive, Charlottesville, VA, 22901 Support (434) 964 4100

### Figure legends

**Figure 1 | Schematic illustration of a hypothetical 3 min fluctuating time course of pain rating.** The variable pain rating is colour-coded in red (low pain) to yellow (high pain). The balanced design ensures a similar amount of phases with rising pain (change - red) and falling pain (change - blue). Phases of stable pain are highlighted in light green. Slider movements (grey) are neither bound to pain amplitude (high, low) nor to change direction (rising, falling).

**Figure 2 | Connectivity pattern for the encoding of pain intensity across all CBP patients (AMP).** Dashed lines indicate negative relationships with pain intensity; solid lines indicate positive relationships with pain intensity (link).

**Figure 3 | Connectivity pattern for the encoding of the change of pain intensity across all CBP patients (SLP).** Dashed lines indicate negative relationships with rising pain intensity; solid lines indicate positive relationships with rising pain intensity (link).

**Figure 4 | Confusion matrices for the encoding of AMP (left) and SLP (right) for all CBP subjects.** Confusion matrices for AMP and SLP for all 408 region pairs given in t-values: 1-180: left hemisphere, 181-360: right hemisphere; 361-381: R: additional regions; 382-408: C: Cerebellum. AMP shows ~70% negative and ~30% positive significant t-values; the confusion matrix for SLP shows even more negative (~74%) than positive (~26%) ones.

**Figure 5 | Connectivity pattern for the encoding of pain intensity changes across all CM patients (SLP).** Dashed lines indicate negative relationships with rising pain intensity; solid lines indicate positive relationships with rising pain intensity (link).

**Figure 6 | Connectivity pattern for the encoding of pain intensity across all CM patients (AMP).** Dashed lines indicate negative relationships with rising pain intensity; solid lines indicate positive relationships with rising pain intensity (link).

**Figure 7 | Confusion matrices for the encoding of AMP and SLP for all CM subjects.** Confusion matrices for AMP and SLP for all 408 region pairs given in t-values: 1-180: left hemisphere, 181-360: right hemisphere; 361-381: additional regions (R); 382-408: Cerebellum (C). The confusion matrix for AMP shows 25% positive significant t-values, and 75% negative ones, compared to SLP which shows more positive (~58%) than negative (~42%) ones.

For Peer Review

### Abbreviated Summary

Mayr and colleagues investigated how the ongoing intensity of chronic pain is reflected by the evolving cortical connectivity. At group level, we find decreased connectivity for rising and high pain. Group results do not reflect the connectivity patterns of individual patients.

For Peer Review

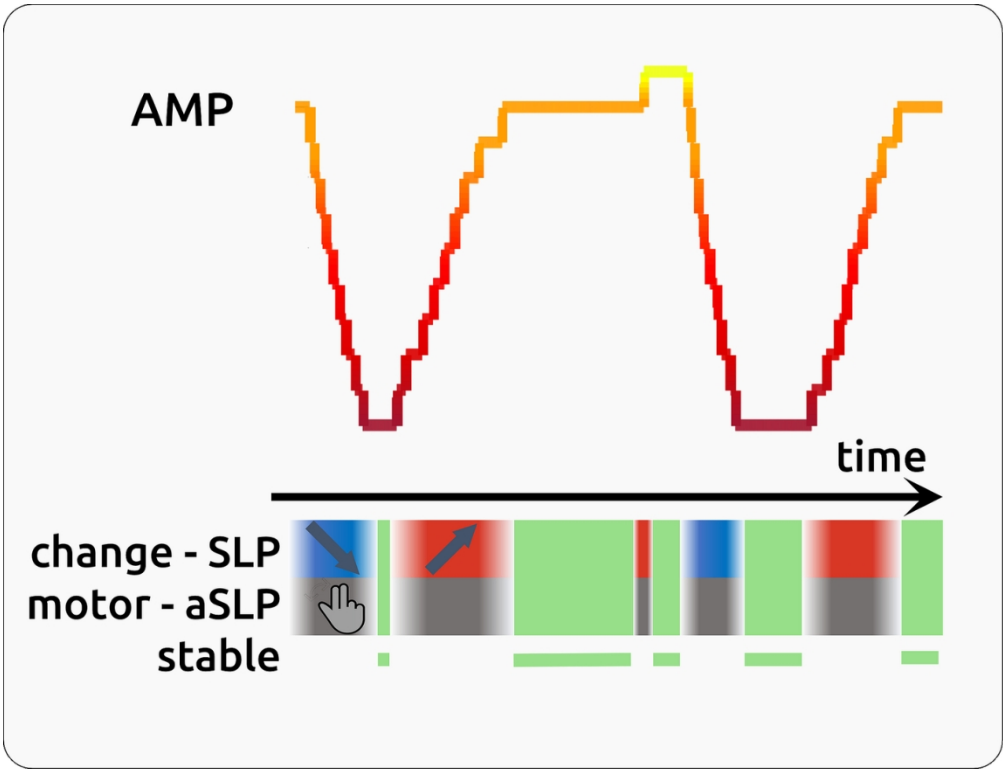

Figure 1 | Schematic illustration of a hypothetical 3 min fluctuating time course of pain rating. The variable pain rating is colour-coded in red (low pain) to yellow (high pain). The balanced design ensures a similar amount of phases with rising pain (change - red) and falling pain (change - blue). Phases of stable pain are highlighted in light green. Slider movements (grey) are neither bound to pain amplitude (high, low) nor to change direction (rising, falling).

119x91mm (300 x 300 DPI)

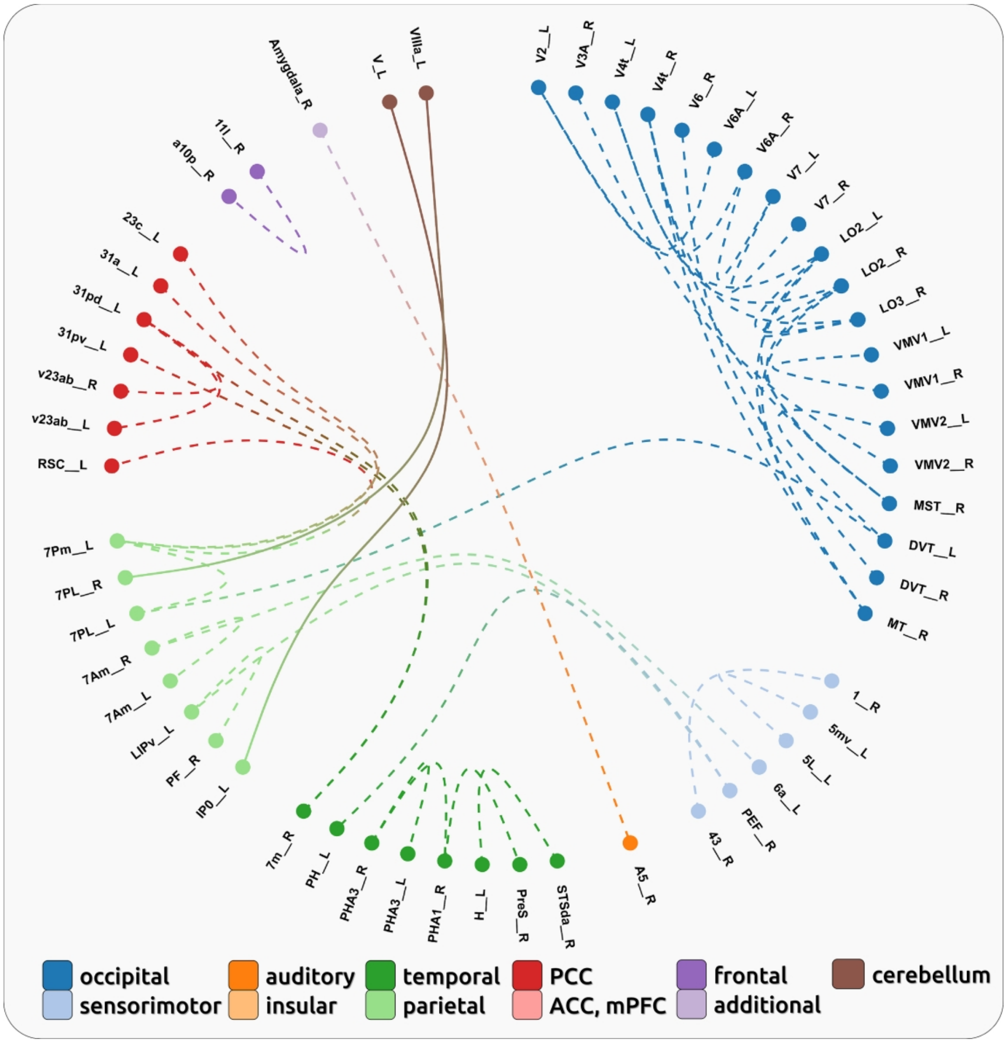

Figure 2 | Connectivity pattern for the encoding of pain intensity across all CBP patients (AMP). Dashed lines indicate negative relationships with pain intensity; solid lines indicate positive relationships with pain intensity (link).

149x155mm (600 x 600 DPI)

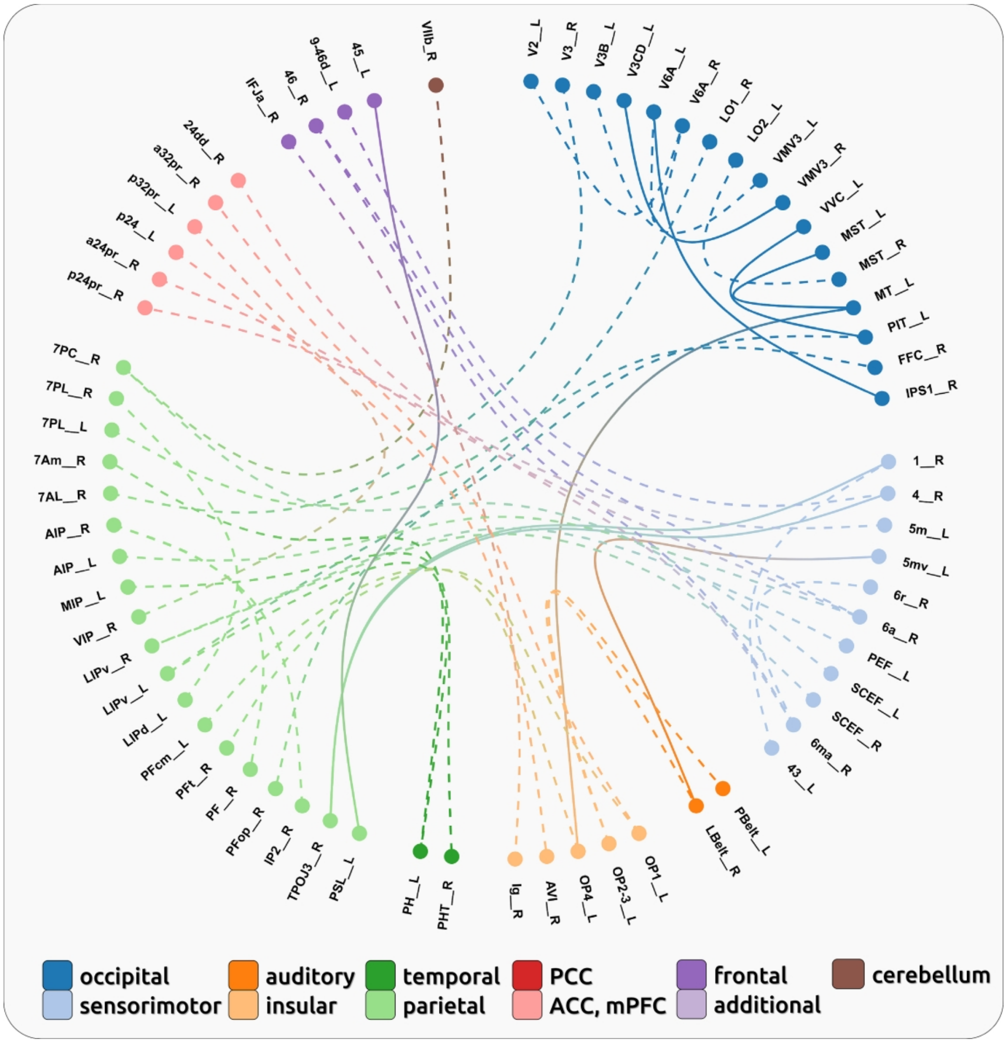

Figure 3 | Connectivity pattern for the encoding of the change of pain intensity across all CBP patients (SLP). Dashed lines indicate negative relationships with rising pain intensity; solid lines indicate positive relationships with rising pain intensity (link).

149x155mm (600 x 600 DPI)

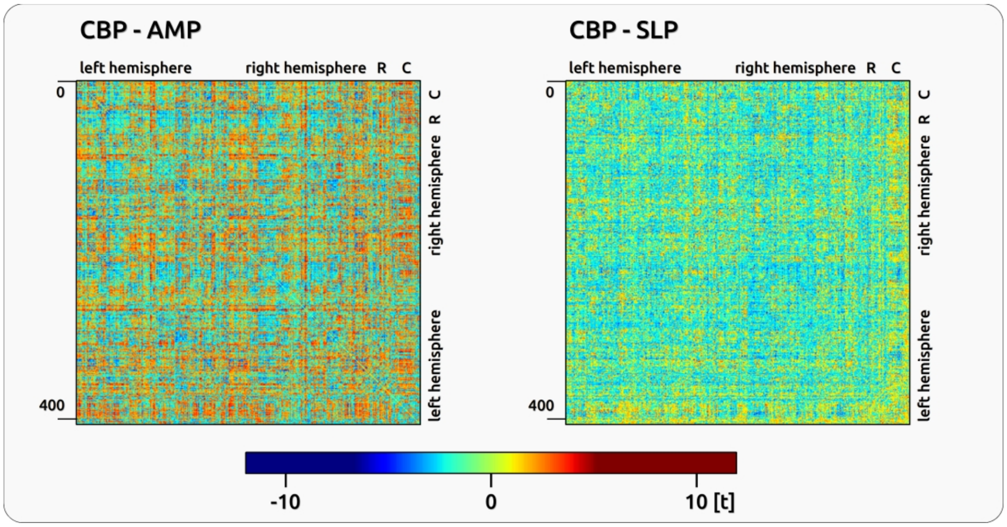

Figure 4 | Confusion matrices for the encoding of AMP (left) and SLP (right) for all CBP subjects. Confusion matrices for AMP and SLP for all 408 region pairs given in t-values: 1-180: left hemisphere, 181-360: right hemisphere; 361-381: R: additional regions; 382-408: C: Cerebellum. AMP shows ~70% negative and ~30% positive significant t-values; the confusion matrix for SLP shows even more negative (~74%) than positive (~26%) ones.

149x77mm (600 x 600 DPI)

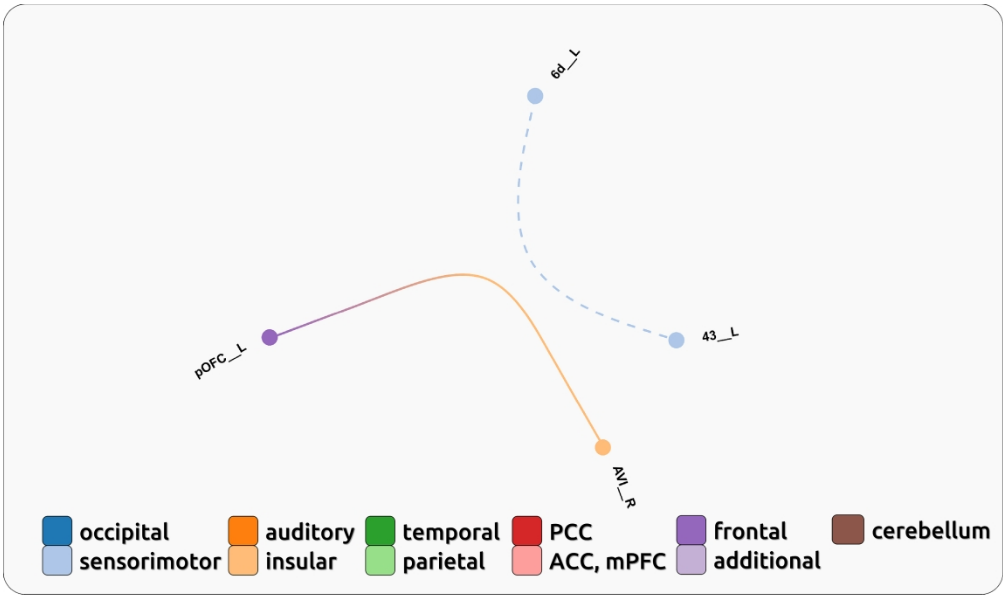

Figure 5 | Connectivity pattern for the encoding of pain intensity changes across all CM patients (SLP). Dashed lines indicate negative relationships with rising pain intensity; solid lines indicate positive relationships with rising pain intensity (link).

149x88mm (600 x 600 DPI)

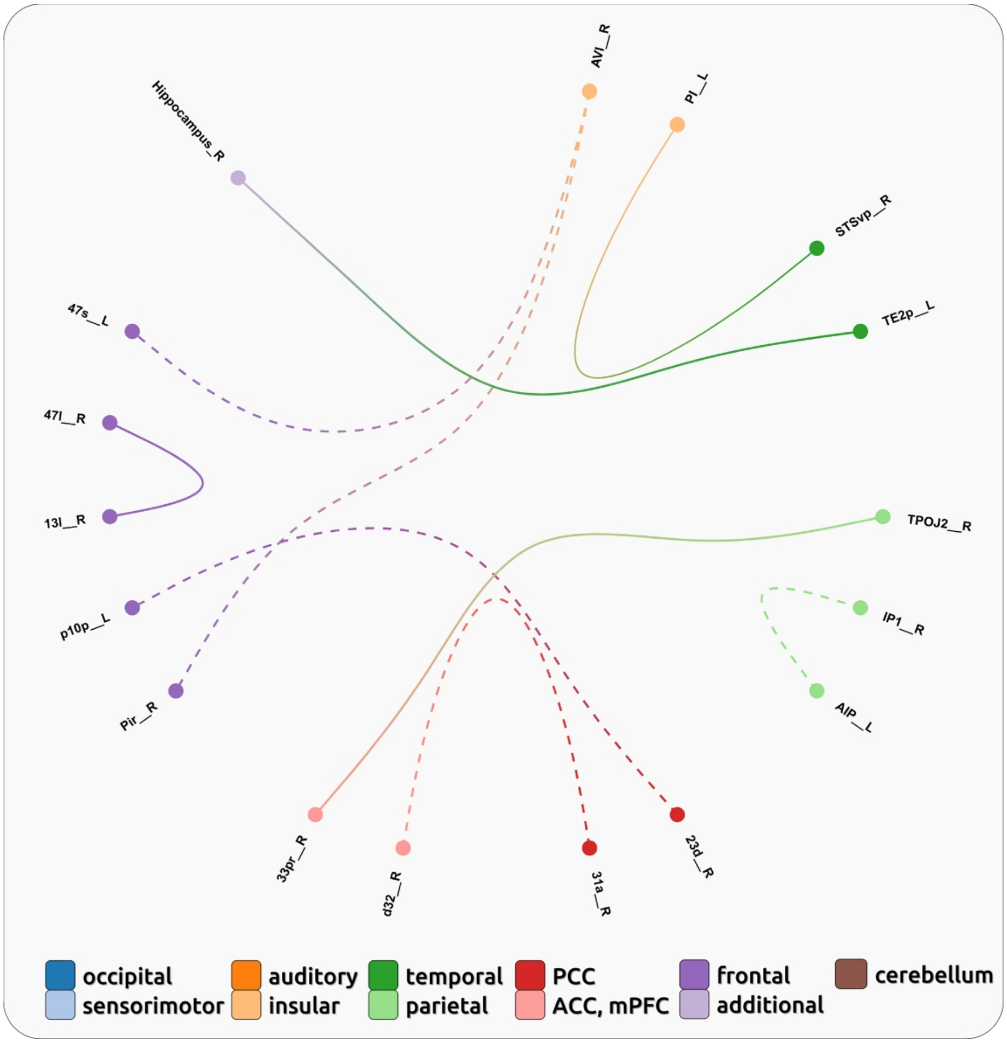

Figure 6 | Connectivity pattern for the encoding of pain intensity across all CM patients (AMP). Dashed lines indicate negative relationships with rising pain intensity; solid lines indicate positive relationships with rising pain intensity (link).

149x155mm (600 x 600 DPI)

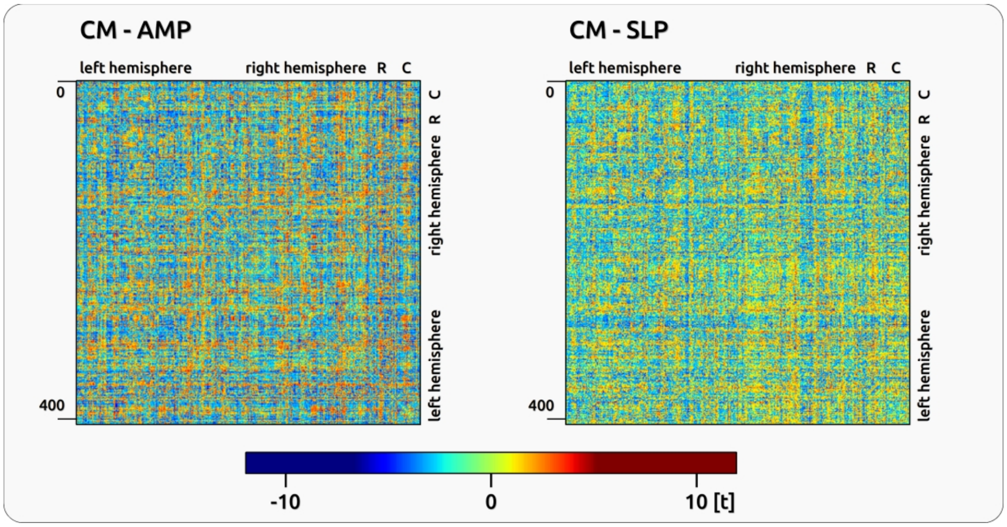

Figure 7 | Confusion matrices for the encoding of AMP and SLP for all CM subjects. Confusion matrices for AMP and SLP for all 408 region pairs given in t-values: 1-180: left hemisphere, 181-360: right hemisphere; 361-381: additional regions (R); 382-408: Cerebellum (C). The confusion matrix for AMP shows 25% positive significant t-values, and 75% negative ones, compared to SLP which shows more positive (~58%) than negative (~42%) ones.

149x77mm (600 x 600 DPI)

**Supplementary Table 1 | Characteristics of CBP patients reported in questionnaires.**

| # | m/f | age (years) | pain duration (years) | pain location | pain medication | pain intensity | PCS | d/a/s |
|---|---|---|---|---|---|---|---|---|
| 1 | f | 52 | 32 | thoracic | none | 4 | 14 | 1/4/11 |
| 2 | m | 64 | 2 | lumbar | none | 3 | 22 | 3/5/7 |
| 3 | f | 39 | 8 | lumbar | Ibuprofen 600 mg (10-12x/month)) | 4 | 32 | 7/9/16 |
| 4 | f | 41 | 18 | lumbar | Fluoxetine 20 mg (daily), Paracetamol 500 mg (2-3x/month), Orphenadrine 100mg (3x/month) | 3 | 33 | 6/7/11 |
| 5 | f | 60 | 14 | thoracic | Diclofenac 69,82 mg (1x/month) | 4 | 14 | 2/5/6 |
| 6 | f | 39 | 3 | lumbar | Paracetamol 500mg (2-3x/month), Ibuprofen 400mg (2x/month) | 4 | 13 | 2/1/1 |
| 7 | f | 48 | 5 | cervical | Ibuprofen 400mg (4x/month) | 5 | 12 | 4/0/6 |
| 8 | f | 52 | 11 | thoracic & lumbar | Ibuprofen 400mg (5x/month), Metamizole 500mg (2x/month) | 4 | 7 | 1/2/5 |
| 9 | f | 31 | 6 | lumbar | Ibuprofen 800mg (2-3x/month), Metamizole 1000mg (2-3x/month) | 5 | 1 | 0/0/1 |
| 10 | m | 26 | 4 | lumbar | none | 5 | 9 | 0/3/3 |
| 11 | f | 55 | 16 | thoracic & lumbar | Cannabis drops (15x/month) | 10 | 11 | 5/2/6 |
| 12 | m | 65 | 8 | lumbar | Ibuprofen 600mg (10-15x/month) | 4 | 10 | 2/2/3 |
| 13 | f | 31 | 1 | cervical & thoracic | Ibuprofen 400mg (18x/month) | 3 | 25 | 9/6/14 |
| 14 | m | 32 | 7 | thoracic and lumbar | Ibuprofen 600mg (6x/month), Tramadol 100mg (10x/month) | 5 | 22 | 7/3/5 |
| 15 | f | 26 | 10 | lumbar | Ibuprofen 400mg (1x/month) | 4 | 18 | 6/3/8 |
| 16 | f | 55 | 16 | cervical & thoracic | Metamizole 500mg (2-3x/month) | 5 | 2 | 4/0/4 |
| 17 | f | 56 | 15 | lumbar | none | 7 | 36 | 4/4/13 |
| 18 | f | 42 | 11 | thoracic & lumbar | none | 5 | 10 | 1/2/6 |
| 19 | f | 30 | 3 | thoracic | Ibuprofen 400mg (3x/month) | 4 | 27 | 2/4/8 |
| 20 | f | 43 | 10 | cervical & lumbar | Ibuprofen 400mg (5x/month) | 7 | 22 | 5/4/6 |

m/f: male/female; PCS: pain catastrophizing scale; d/a/s: depression/anxiety/stress. The cutoff for depression and stress is 10, for anxiety 6, and for the PCS 30.

**Supplementary Table 2 | Characteristics of CM patients reported in questionnaires.**

| # | m/f | age (years) | pain duration (years) | pain medication | pain intensity | PCS | d/a/s |
|---|---|---|---|---|---|---|---|
| 1 | f | 61 | 50 | Sumatriptan 100mg (20x/month) | 7 | 15 | 0/10/9 |
| 2 | f | 27 | 7 | Metamizole 500mg (2-3x/month), Sumatriptan 50mg (1x/month) | 4 | 5 | 0/0/2 |
| 3 | f | 50 | 35 | Sumatriptan 100mg (5-7x/month) | 4 | 24 | 5/0/6 |
| 4 | f | 27 | 8 | Zolmitriptan 20mg (2x/month), Ibuprofen 600mg (7x/month) | 7 | 37 | 1/1/8 |
| 5 | m | 49 | 30 | Ibuprofen 600mg (7-8x/month), Metamizole 500mg (3-4x/month), Paracetamol 500mg (5-6x/month) | 4 | 3 | 0/6/1 |
| 6 | f | 52 | 30 | Ibuprofen 400mg (6x/month), Paracetamol 1000mg (2x/month) | 5 | 11 | 5/10/12 |
| 7 | f | 32 | 15 | Zolmitriptan 5mg (8x/month), Naproxen 500mg (15x/month), Acetylsalicylic Acid (ASA) 250mg (4x/month), Paracetamol 200mg (4x/month), Caffeine 50mg (4x/month) | 4 | 31 | 5/7/7 |
| 8 | f | 21 | 7 | Sumatriptan 50mg (1x/month) | 4 | 10 | 2/1/0 |
| 9 | f | 19 | 7 | none | 4 | 35 | 10/2/6 |
| 10 | f | 46 | 13 | Ibuprofen 800mg (8-10x/month) | 6 | 31 | 7/10/15 |
| 11 | f | 27 | 13 | Triptan (2-3x/month), ASA 250mg (20-25x/month), Paracetamol 250mg (20-25x/month), Caffeine 50mg (20-25x/month) | 4 | 13 | 6/1/10 |
| 12 | m | 53 | 15 | Ibuprofen 600mg (10x/month) | 6 | 20 | 7/2/4 |
| 13 | f | 30 | 6 | Ibuprofen 400mg (4x/month) | 4 | 24 | 1/1/5 |
| 14 | f | 21 | 7 | Ibuprofen 600mg (4-8x/month), Paracetamol 500mg (4x/month), Zolmitriptan 5mg (1-2x/month) | 3 | 15 | 2/0/5 |
| 15 | f | 23 | 8 | Ibuprofen 600mg (2-3x/month) | 7 | 24 | 2/0/3 |
| 16 | f | 28 | 7 | Ibuprofen 500mg (10x/month) | 7 | 11 | 0/1/1 |
| 17 | f | 25 | 5 | Ibuprofen 600mg (5x/month), Zolmitriptan 5mg (1x/month) | 4 | 21 | 5/5/9 |
| 18 | f | 33 | 20 | Paracetamol 500mg (3x/month) | 5 | 32 | 4/0/4 |
| 19 | f | 21 | 9 | Ibuprofen 400mg (6-10x/month), Rizatriptan 10mg (2x/month) | 5 | 30 | 1/1/2 |
| 20 | f | 43 | 10 | Ibuprofen 400mg (20x/month), Paracetamol 325 mg (8-10x/month), Naproxen 100 mg (8-10x/month), Caffeine 50 mg (8-10x/month), Drotaverine hydrochloride 40 mg (8-10x/month), Pheniramine 10 mg (8-10x/month) | 4 | 22 | 2/3/9 |

m/f: male/female; PCS: pain catastrophizing scale; d/a/s: depression/anxiety/stress. The cutoff for depression and stress is 10, for anxiety 6, and for the PCS 30.

**Supplementary Table 3 | MNI coordinates of additional regions.**

| Anatomical subcortical structure (Left or Right) | MNI coordinates | | |
|---|---|---|---|
| | x | y | z |
| Accumbens L | -7 | 9 | -8 |
| Accumbens R | 8 | 11 | -8 |
| Amygdala L | -23 | -5 | -20 |
| Amygdala R | 24 | -4 | -20 |
| Brain Stem | 0 | -30 | -33 |
| Hippocampus L | -25 | -23 | -14 |
| Hippocampus R | 26 | -22 | -14 |
| Pallidum L | -19 | -4 | -2 |
| Pallidum R | 20 | -4 | -2 |
| Thalamus L | -11 | -19 | 6 |
| Thalamus R | 12 | -18 | 6 |
| Hypothalamus | 6 | 0 | -12 |
| Hypothalamus #2 | 6 | -6 | -12 |
| Locus Coeruleus | 3 | -38 | -23 |
| Ncl Caudatus L | -13 | 9 | 10 |
| Putamen L | -25 | 0 | 1 |
| PAG | 1 | -34 | -8 |
| Pons | 4 | -20 | -20 |
| Ncl Caudatus R | 14 | 10 | 11 |
| Putamen R | 26 | 2 | 0 |
| Spinal Sphere | 6 | -40 | -46 |

**Supplementary Table 4 | MNI coordinates of additional Cerebellum regions.**

| Anatomical Cerebellum structure (Left or Right) | MNI coordinates | | |
|---|---|---|---|
| | x | y | z |
| ItoIV L | -7 | -44 | -17 |
| ItoIV R | 10 | -43 | -18 |
| V L | -13 | -50 | -19 |
| V R | 14 | -51 | -19 |
| VI L | -23 | -59 | -25 |
| VermisVI | 1 | -71 | -21 |

| | | | |
|---|---|---|---|
| VI R | 24 | -58 | -25 |
| CrusI L | -36 | -68 | -32 |
| CrusI R | 38 | -68 | -32 |
| CrusII L | -26 | -75 | -42 |
| VermisCrusII | 0 | -75 | -31 |
| CrusII R | 26 | -76 | -41 |
| VIIb L | -26 | -66 | -51 |
| VermisVIIb | 0 | -68 | -31 |
| VIIb R | 28 | -65 | -50 |
| VIIIa L | -24 | -57 | -53 |
| VermisVIIIa | 0 | -67 | -38 |
| VIIIa R | 26 | -58 | -53 |
| VIIIb L | -17 | -50 | -55 |
| VermisVIIIb | 0 | -63 | -42 |
| VIIIb R | 18 | -51 | -55 |
| IX L | -7 | -53 | -48 |
| VermisIX | 0 | -56 | -37 |
| IX R | 7 | -53 | -48 |
| X L | -21 | -37 | -45 |
| VermisX | 1 | -48 | -35 |
| X R | 22 | -37 | -46 |

**Supplementary Table 5 Regions with more than or equal to two connections encoding the pain intensity across all CBP patients (AMP).**

| | # of connections | Area Description | Area name | L | R |
|---|---|---|---|---|---|
| 1 | 4 | Medial Area 7P | 7Pm | L | |
| 2 | 4 | Area Lateral Occipital 2 | LO2 | L | |
| 3 | 4 | Area Lateral Occipital 2 | LO2 | | R |
| 4 | 3 | Area 31pd | 31pd | L | |
| 5 | 3 | Medial Superior Temporal Area | MST | | R |
| 6 | 3 | Area Lateral Occipital 3 | LO3 | | R |
| 7 | 3 | Middle Temporal Area | MT | | R |
| 8 | 3 | Area V4t | V4t | | R |
| 9 | 2 | Area 7m | 7m | | R |
| 10 | 2 | Dorsal Transitional Visual Area | DVT | L | |
| 11 | 2 | Lateral Area 7P | 7PL | L | |
| 12 | 2 | Medial Area 7A | 7Am | | R |
| 13 | 2 | Area Lateral IntraParietal ventral | LIPv | L | |
| 14 | 2 | Premotor Eye Field | PEF | | R |
| 15 | 2 | ParaHippocampal Area 1 | PHA1 | | R |
| 16 | 2 | ParaHippocampal Area 3 | PHA3 | | R |
| 17 | 2 | Auditory 5 Complex | A5 | | R |
| 18 | 2 | Seventh Visual Area | V7 | L | |
| 19 | 2 | Area V4t | V4t | L | |
| 20 | 2 | Area V6A | V6A | | R |

**Supplementary Table 6 | Regions with more than or equal to two connections encoding the change in pain intensity across all CBP patients (SLP).**

| | # of connections | Area Description | Area name | L | R |
|---|---|---|---|---|---|
| 1 | 3 | Area 6m anterior | 6ma | | R |
| 2 | 3 | Area 6 anterior | 6a | | R |
| 3 | 3 | Area V6A | V6A | | R |
| 4 | 3 | Area V6A | V6A | L | |
| 5 | 2 | Area 1 ( primary somatosensory cortex) | 1 | | R |
| 6 | 2 | Primary Motor Cortex | 4 | | R |
| 7 | 2 | Middle Temporal Area | MT | L | |
| 8 | 2 | Posterior InferoTemporal | PIT | L | |
| 9 | 2 | Area OP1/SII | OP1 | L | |
| 10 | 2 | Area OP2-3/VS | OP2-3 | L | |
| 11 | 2 | Area OP4/PV | OP4 | L | |
| 12 | 2 | Area TemporoParietoOccipital Junction 3 | TPOJ3 | | R |
| 13 | 2 | Lateral Belt Complex | LBelt | | R |
| 14 | 2 | Area PH | PH | L | |
| 15 | 2 | Area Lateral IntraParietal ventral | LIPv | | R |
| 16 | 2 | Area Lateral IntraParietal ventral | LIPv | L | |
| 17 | 2 | Anterior IntraParietal Area | AIP | | R |
| 18 | 2 | Area 7PC | 7PC | | R |
| 19 | 2 | Area 46 | 46 | | R |

**Supplementary Table 7 | Correlation between individual maps and the group maps indicated by Kendall's tau.**

| | CBP | | | CM | | |
|---|---|---|---|---|---|---|
| | **AMP** | **SLP** | **aSLP** | **AMP** | **SLP** | **aSLP** |
| 1 | 0.053 | 0.065 | 0.095 | 0.12 | 0.070 | 0.20 |
| 2 | 0.20 | 0.027 | 0.081 | 0.093 | 0.061 | 0.27 |
| 3 | 0.055 | 0.030 | 0.097 | 0.017 | 0.0072 | -0.050 |
| 4 | 0.10 | 0.0044 | -0.0084 | 0.022 | 0.050 | 0.12 |
| 5 | 0.043 | -0.018 | -0.027 | 0.076 | 0.0052 | 0.12 |
| 6 | 0.12 | -0.00032 | 0.22 | 0.071 | 0.099 | 0.0046 |
| 7 | 0.085 | 0.072 | 0.075 | 0.12 | 0.063 | 0.084 |
| 8 | 0.14 | 0.027 | 0.14 | 0.096 | 0.0064 | 0.069 |
| 9 | 0.039 | 0.0024 | 0.080 | 0.058 | 0.052 | 0.11 |
| 10 | 0.13 | 0.047 | 0.075 | 0.080 | 0.036 | -0.026 |
| 11 | 0.086 | 0.051 | 0.22 | 0.085 | 0.073 | 0.079 |
| 12 | 0.12 | 0.0015 | 0.084 | 0.15 | 0.050 | 0.16 |
| 13 | 0.068 | 0.11 | 0.061 | 0.040 | 0.027 | 0.13 |
| 14 | 0.099 | 0.034 | 0.082 | 0.16 | 0.098 | 0.089 |
| 15 | -0.0079 | 0.12 | 0.022 | 0.033 | 0.054 | 0.076 |
| 16 | 0.089 | 0.0027 | 0.13 | 0.095 | -0.012 | 0.19 |
| 17 | 0.0036 | 0.034 | 0.022 | 0.053 | 0.085 | 0.14 |
| 18 | 0.10 | -0.020 | 0.072 | 0.036 | 0.087 | 0.12 |
| 19 | 0.052 | -0.019 | 0.16 | 0.016 | 0.0085 | 0.075 |
| 20 | 0.14 | 0.068 | 0.040 | 0.056 | 0.035 | -0.052 |

**Supplementary Table 8 | Number of significant connections for every single subject for CBP and CM for the encoding of AMP and SLP.**

| | # of significant connections for CBP | | # of significant connections for CM | |
|---|---|---|---|---|
| | **AMP** | **SLP** | **AMP** | **SLP** |
| 1 | 106 | 106 | 11 | 7 |
| 2 | 65 | 13 | 70 | 29 |
| 3 | 44 | 188 | 24 | 29 |
| 4 | 1240 | 55 | 20 | 33 |
| 5 | 54 | 5 | 59 | 10 |
| 6 | 109 | 462 | 41 | 1 |
| 7 | 95 | 2284 | 24 | 59 |
| 8 | 15 | 26 | 255 | 460 |
| 9 | 131 | 0 | 36 | 2 |
| 10 | 20 | 55 | 52 | 14 |
| 11 | 2 | 4 | 4 | 52 |
| 12 | 3 | 4 | 39 | 163 |
| 13 | 158 | 2716 | 128 | 32 |
| 14 | 78 | 8 | 17 | 17 |
| 15 | 704 | 1437 | 12 | 5889 |
| 16 | 7 | 19 | 5 | 66 |
| 17 | 164 | 4 | 197 | 371 |
| 18 | 14 | 0 | 151 | 2888 |
| 19 | 92 | 65 | 112 | 583 |
| 20 | 26 | 5857 | 508 | 73 |

**Supplementary Figure 1 | Accepted and rejected pain ratings.**

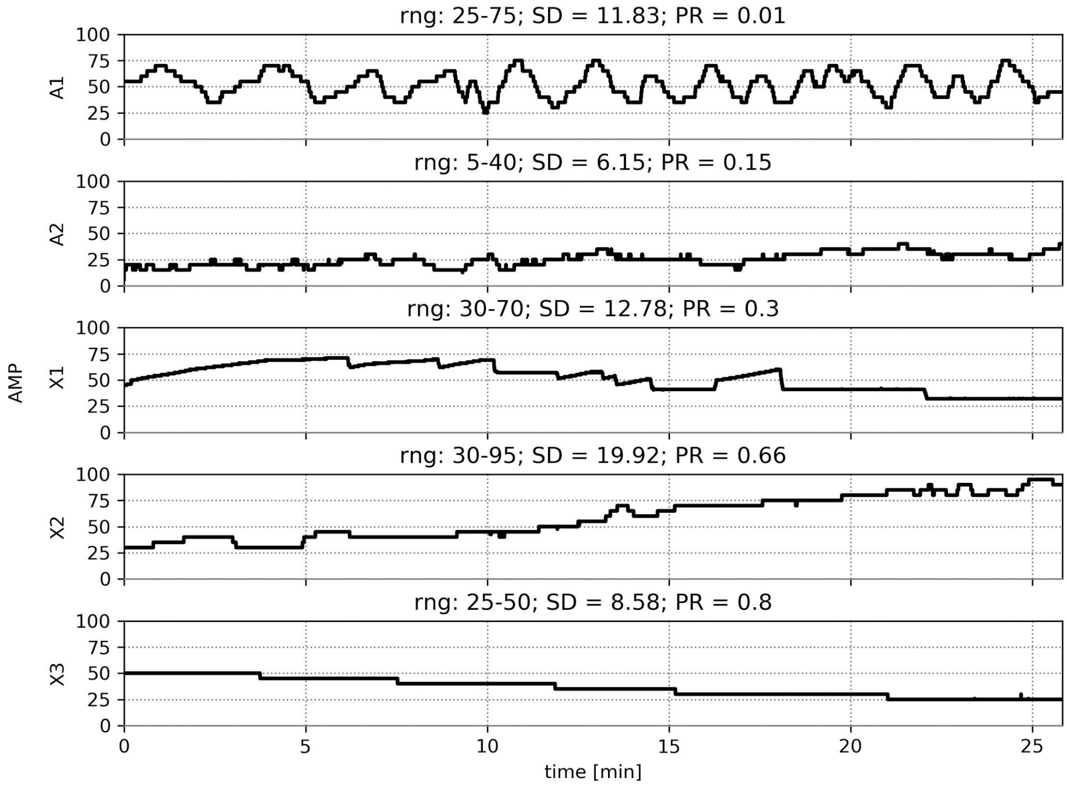

Accepted and rejected pain ratings based on the parameter PR. Rating A1 represents an excellent rating with a very low PR value of 0.01, indicating high variance and no overall drift throughout the experiment. Rating A2 represents a pain rating with a moderate PR value of 0.15, whereas X1 represents an excluded rating with a PR slightly higher than the threshold PR ≥ 0.25. X2 was excluded due to a steady increase in the pain ratings over the course of the experiment. X3 was excluded due to very low variability in ratings after high-pass filtering. Ratings A1 and A2 were accepted, whereas recordings X1, X2 and X3 were excluded from the analysis.

The ratings of each patient's pain was measured with the parameter PR defined as follows, with "$\Delta pain_{ud}/\Delta time$" being the slope of the regression line of the unfiltered data (ud) and $\sigma_{fd}$ being the sample standard deviation of the filtered data (fd):

$$(1)\ PR = \left| \frac{\Delta pain_{ud} / \Delta time}{\sigma_{fd}} \right|$$

The parameter PR is constructed in a way that its minimisation is desirable. The numerator describes how much the prerequisite is violated by fitting a least squares line across the rating time course. This violation can be compensated if the variance of pain ratings is at least four times higher than the slope of the regression of the least squares line across the entire time course of the pain ratings. The standard deviation of the filtered data expressed in the denominator gives a measure of the desired fluctuation of the pain ratings but is stripped from a potential trend throughout the experiment. A minimisation of the quotient is given either by minimising the numerator corresponding to a small increase of the overall pain ratings over the whole experiment (small slope), or by maximising the denominator corresponding to a large variability in pain ratings across the rating task. Minor overall rising in pain intensity over the whole time of the experiment could be compensated by a greater variance of ratings; small fluctuation of pain intensity would only be accepted in cases of minor pain rating trends across the entire experiment. Recordings showing PR values of ≥0.25 were rejected from the analysis or repeated if possible. We excluded five participants and repeated three sessions. The threshold of the PR value was chosen based on theoretical considerations (effect of order, data filtering) as well as on a careful inspection of the data. Please note that subjects with a *high PR* value are *not suitable* for a continuous pain rating design for two reasons. Firstly, steadily changing ratings will cause an effect of order. Secondly, the main changes of brain activity (see X2 and X3 in Supplementary Figure 1) will be removed from the data due to the necessary high-pass filtering. There is no literature we could have relied on in this matter.

**Supplementary Figure 2 | Time courses of pain ratings for CBP patients.**

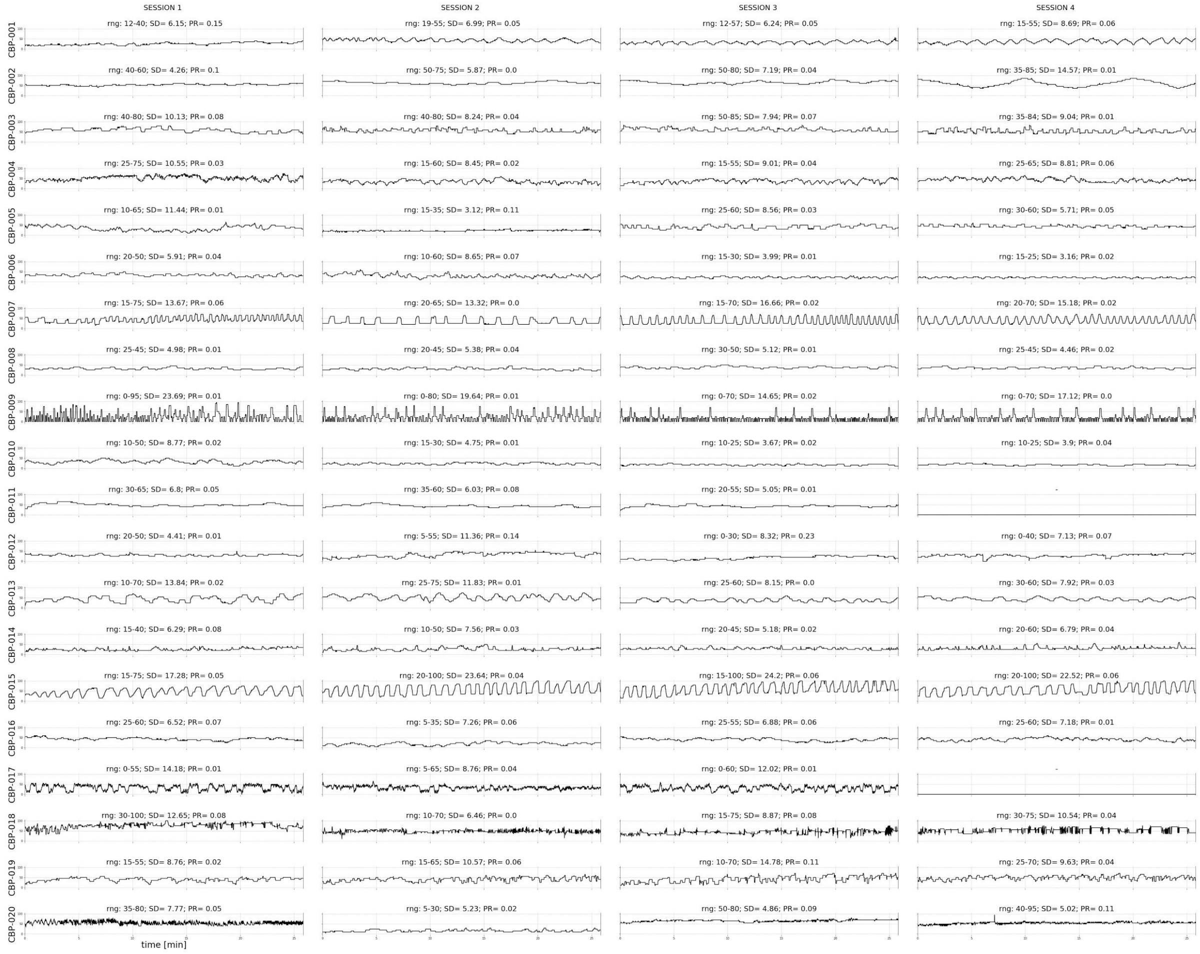

Rated pain intensity (AMP) between 0 and 100 is plotted against time (approximately 25 min total). For all patients (except CBP-011 and CBP-017) four sessions were recorded successfully. The range (rng) of the pain intensity rating and the standard deviation (SD) over the whole time series are given. The ratings of each subject's pain was measured with the parameter PR to ensure a large variability in pain ratings, as well as a small to moderate increase of the overall pain ratings across the rating task.

**Supplementary Figure 3 | Time courses of pain ratings for CM patients.**

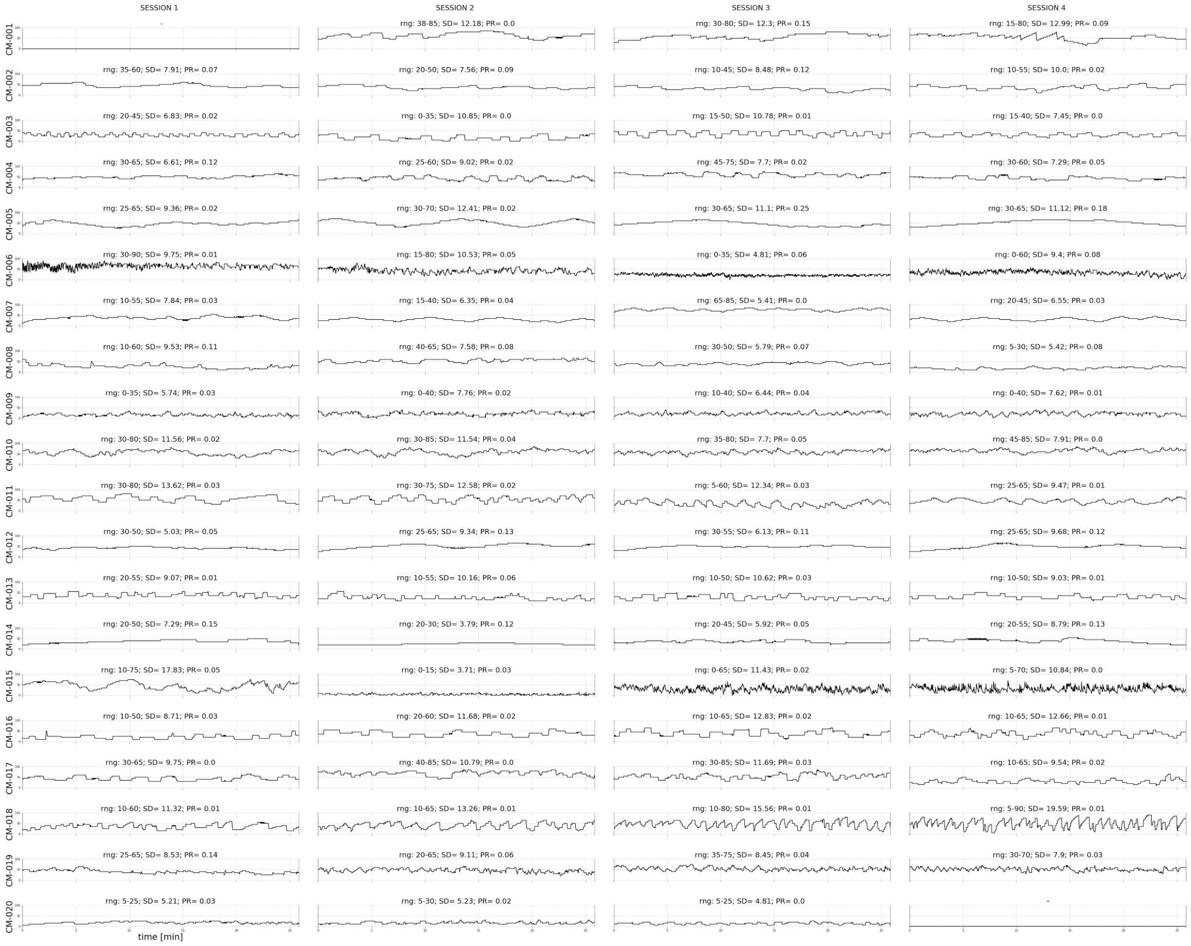

Rated pain intensity (AMP) between 0 and 100 is plotted against time (approximately 25 min total). For all patients (except CM-001 and CM-020), four sessions were recorded successfully. The range (rng) of the pain intensity rating and the standard deviation (SD) over the whole time series are given. The ratings of each subject's pain was measured with the parameter PR to ensure a large variability in pain ratings, as well as a small to moderate increase of the overall pain ratings across the rating task.

**Supplementary Figure 4 | Distribution of shift vectors for CBP and CM connections.**

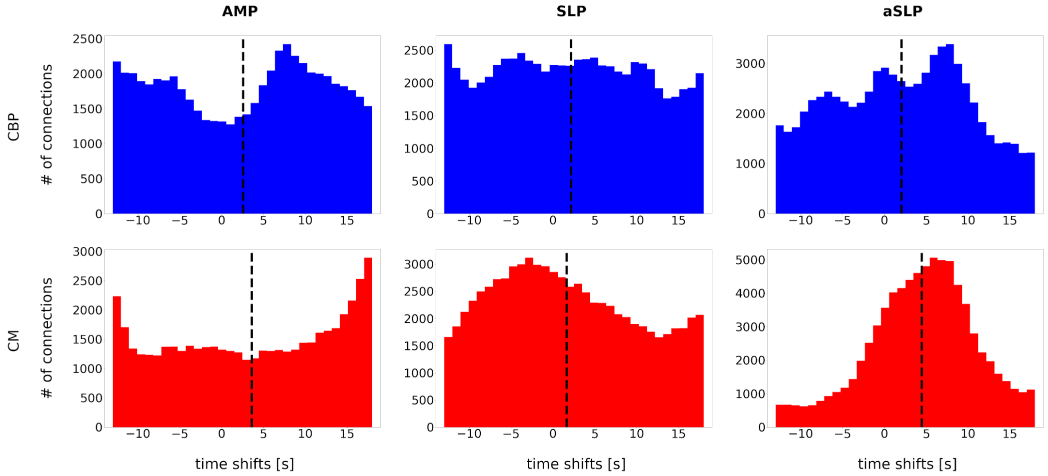

The distribution of shifts was calculated for all connections separately for AMP, SLP, and aSLP for CBP patients (top row, blue) (mean ± SD: AMP (2.6 ± 9.4 s); SLP (2.2 ± 9.1 s); aSLP (2.1 ± 8.3 s)) and for CM patients (bottom row, red) (AMP (3.6 ± 10.0 s); SLP (1.6 ± 8.7 s); aSLP (4.5 ± 6.6 s)) between -14s and 19s. The mean is given as a vertical dashed line for each condition. Standard deviations capture the variability of timing across subjects, all 408 brain regions and four sessions.

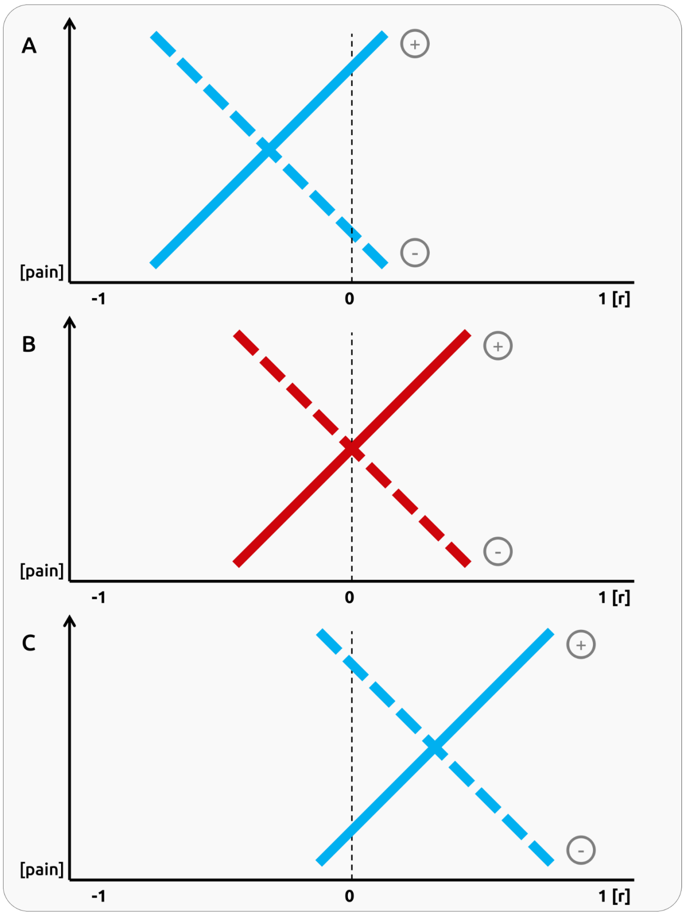

**Supplementary Figure 5 | Potential connectivity schemata.** We considered positive (solid lines) and negative (dashed lines) relationships between functional connectivity and pain intensity ratings (A - C).

*In a first case (A)*, pain-related effects may occur for pairs of brain regions that are largely anti-correlated. A positive relationship with pain intensity would imply a drop of anti-correlation for high pain. Here, low pain states are likely related to suppression effects. This might be the case for "cognitive" brain regions that suppress pain-related insular activity; less activity in these regions and a subsequent disruption of suppression and collapsing anti-correlation would increase pain. For negative pain-related relationships high pain is bound to highly anti-correlated connections; high pain is related to a strong anti-correlation and a suppression effect between brain regions. Higher pain intensities may inhibit frontal brain regions which in turn control behaviour that is involved in the maintenance of pain, e.g. pain-related rumination and hypervigilance on pain. However, we did not find any significant effect for anti-correlated brain regions.

*In a second case (B)*, data would cross the boundary between positive and negative correlation coefficients. We find these cases as non-interpretable, particularly the absence of cortical connectivity for mid-range pain intensities.

*In a third case (C)*, the relationship between pain intensity and functional connectivity occurs for brain regions that are largely correlated. For positive relationships with pain intensity, high-pain states are bound to higher connectivities. The perception of pain may require a crosstalk between two brain regions. A disruption of the crosstalk would lower the intensity of pain. For negative pain-related relationships, high pain states are associated with an decrease of cortical connections. This may occur when one brain region suppresses the processes of the other brain region, e.g. the top-down modulation of the pACC on the PAG. Due to potential noise in the data, we allowed an overlap of 20% towards sections of the data with negative correlation coefficients.

**Supplementary Figure 6 | Individual connectivity patterns.** Circle plots for the encoding of pain intensity (AMP) for all 20 CBP patients (page 1) and 20 CM patients (page 2)

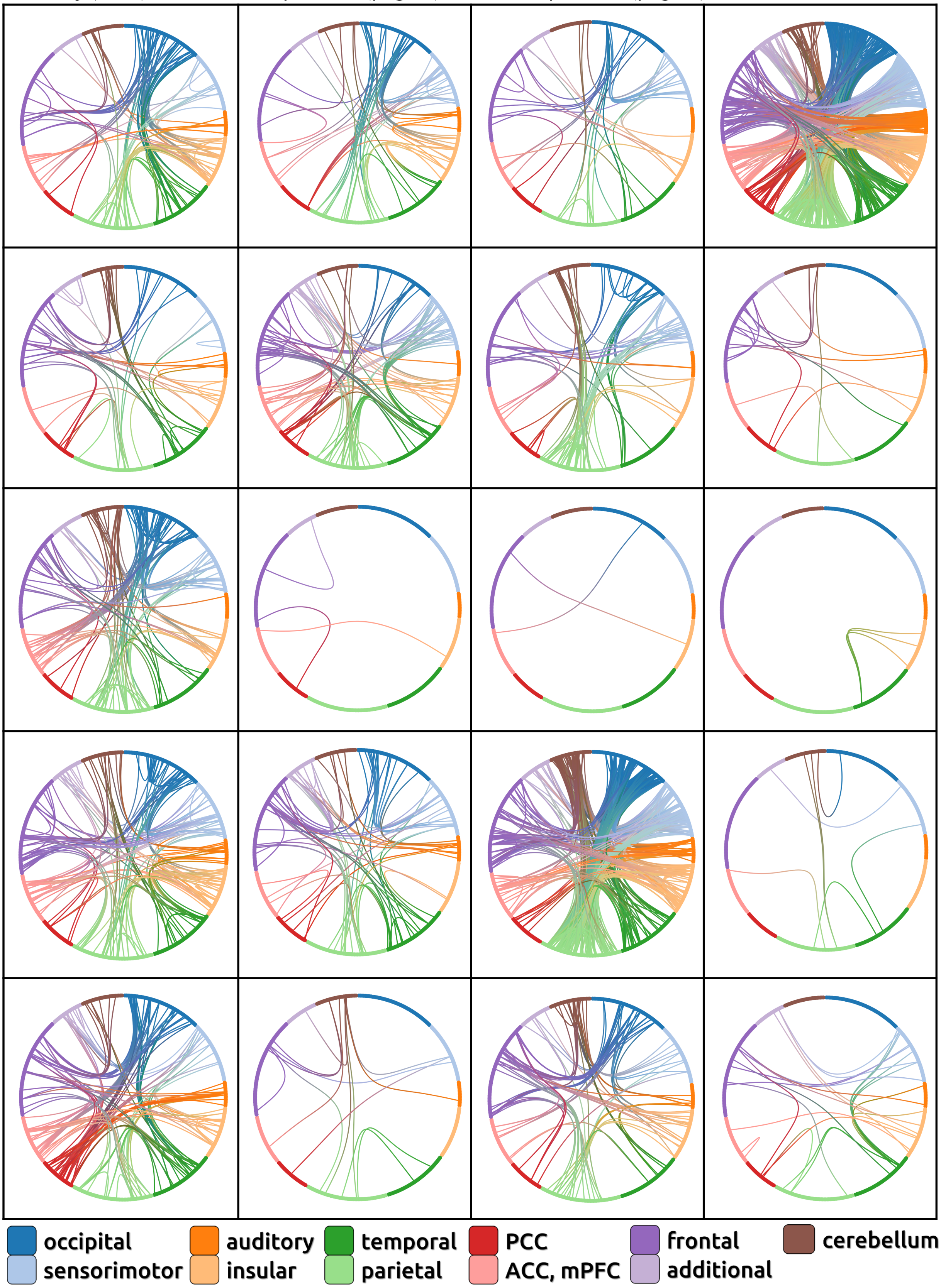

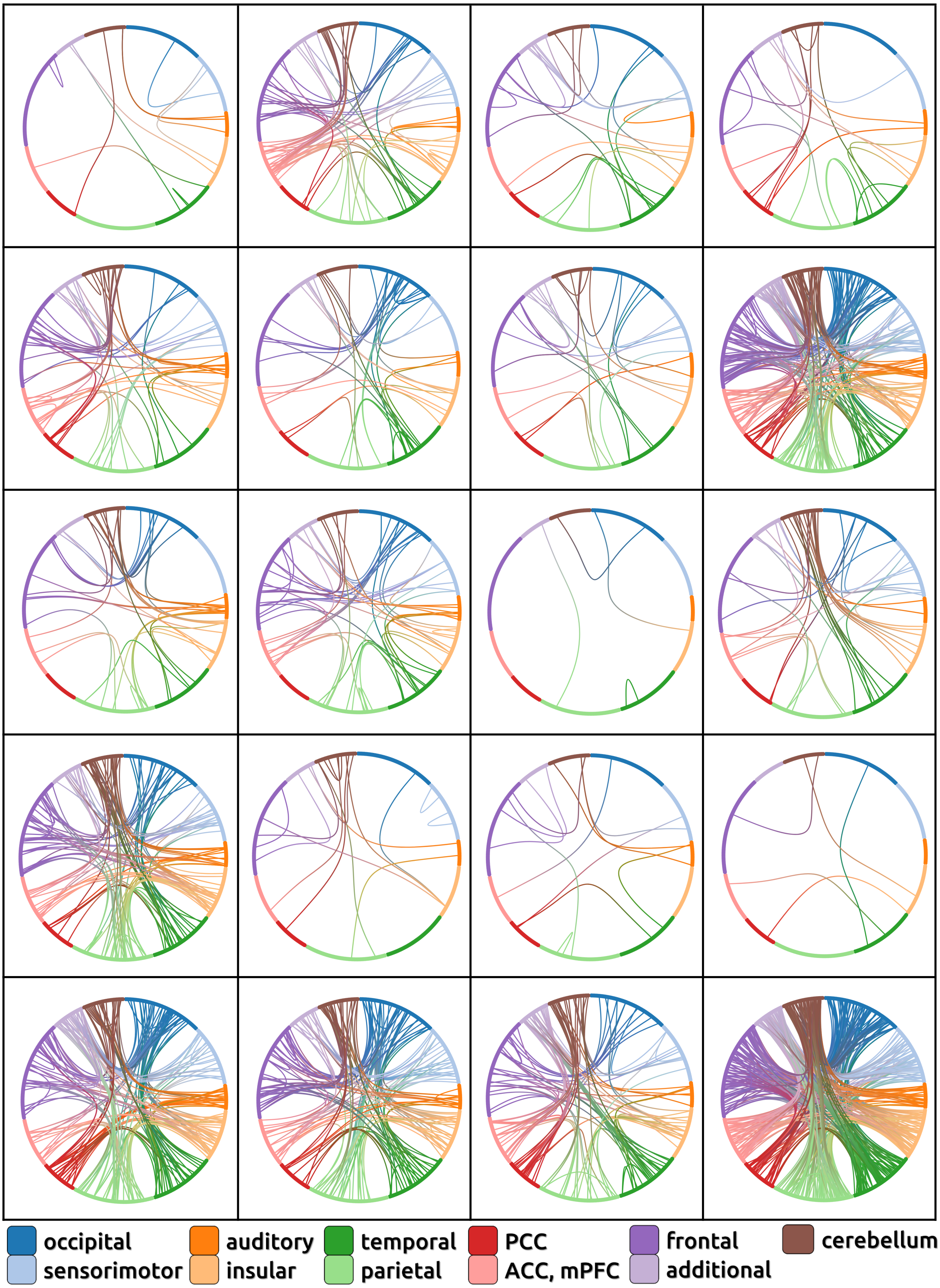
occipital
sensorimotor
auditory
insular
temporal
parietal
PCC
ACC, mPFC
frontal
additional
cerebellum

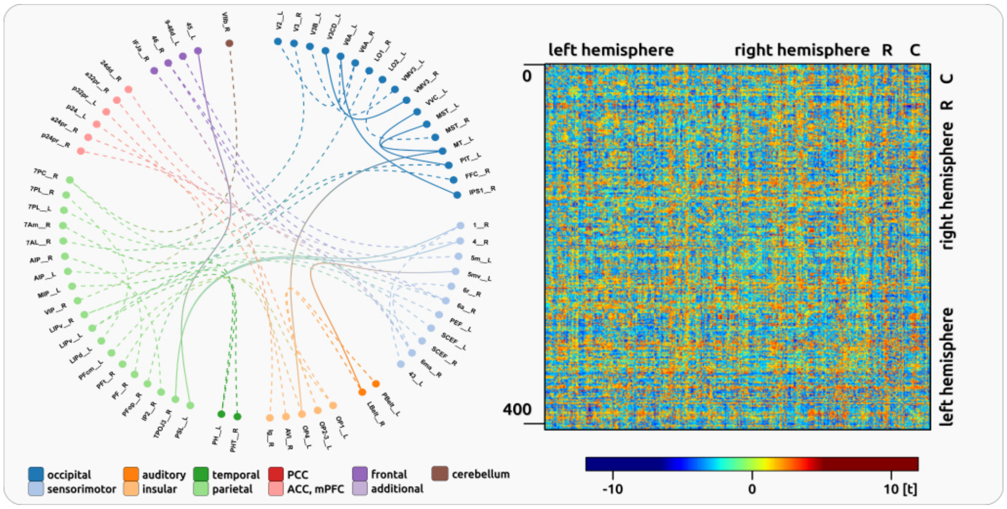

361x180mm (72 x 72 DPI)

### **STROBE** statement: Reporting guidelines checklist for cohort, case-control and cross-sectional studies

| SECTION | ITEM NUMBER | CHECKLIST ITEM | REPORTED ON PAGE NUMBER: |
|---|---|---|---|
| **TITLE AND ABSTRACT** | | | 1 |
| | 1a | Indicate the study's design with a commonly used term in the title or the abstract | 2 |
| | 1b | Provide in the abstract an informative and balanced summary of what was done and what was found | 2 |
| **INTRODUCTION** | | | |
| Background and objectives | 2 | Explain the scientific background and rationale for the investigation being reported | 3-4 |
| | 3 | State specific objectives, including any pre-specified hypotheses | 4 |
| **METHODS** | | | |
| Study design | 4 | Present key elements of study design early in the paper | 5 |
| Setting | 5 | Describe the setting, locations, and relevant dates, including periods of recruitment, exposure, follow-up, and data collection | 5 |
| Participants | 6a | Cohort study—Give the eligibility criteria, and the sources and methods of selection of participants. Describe methods of follow-up<br>Case-control study—Give the eligibility criteria, and the sources and methods of case ascertainment and control selection. Give the rationale for the choice of cases and controls<br>Cross-sectional study—Give the eligibility criteria, and the sources and methods of selection of participants | 5 |
| | 6b | Cohort study—For matched studies, give matching criteria and number of exposed and unexposed<br>Case-control study—For matched studies, give matching criteria and the number of controls per case<br>Variables | n/a |
| Variables | 7 | Clearly define all outcomes, exposures, predictors, potential confounders, and effect modifiers. Give diagnostic criteria, if applicable | 5-7 |

| | | | |
|---|---|---|---|
| Data sources/measurements | 8* | For each variable of interest, give sources of data and details of methods of assessment (measurement). Describe comparability of assessment methods if there is more than one group. | 6 |
| Bias | 9 | Describe any efforts to address potential sources of bias. | 8-9 |
| Study size | 10 | Explain how the study size was arrived at | 5 |
| Quantitative variables | 11 | Explain how quantitative variables were handled in the analyses. If applicable, describe which groupings were chosen and why. | 5-10 |
| Statistical methods | 12a | Describe all statistical methods, including those used to control for confounding | 8-10 |
| | 12b | Describe any methods used to examine subgroups and interactions | 10 |
| | 12c | Explain how missing data were addressed | |
| | 12d | Cohort study—If applicable, explain how loss to follow-up was addressed<br>Case-control study—If applicable, explain how matching of cases and controls was addressed<br>Cross-sectional study—If applicable, describe analytical methods taking account of sampling strategy | n/a |
| | 12e | Describe any sensitivity analyses | n/a |
| **RESULTS** | | | |
| Participants | 13a | Report numbers of individuals at each stage of study—eg numbers potentially eligible, examined for eligibility, confirmed eligible, included in the study, completing follow-up, and analysed | n/a |
| | 13b | Give reasons for non-participation at each stage | n/a |
| | 13c | Consider use of a flow diagram | n/a |
| Descriptive Data | 14a | Give characteristics of study participants (eg demographic, clinical, social) and information on exposures and potential confounders | supplement |
| | 14b | Indicate number of participants with missing data for each variable of interest | n/a |
| | 14c | Cohort study—Summarise follow-up time (eg, average and total amount) | n/a |
| Outcome Data | 15* | Cohort study—Report numbers of outcome events or summary measures over time<br>Case-control study—Report numbers in each exposure category, or summary measures of exposure<br>Cross-sectional study—Report numbers of outcome events or summary measures | n/a |

BRAIN A JOURNAL OF NEUROLOGY
ScholarOne, 375 Greenbrier Drive, Charlottesville, VA, 22901 Support (434) 964 4100

| | | | |
|---|---|---|---|
| Main Results | 16a | Give unadjusted estimates and, if applicable, confounder-adjusted estimates and their precision (e.g. 95% confidence interval). Make clear which confounders were adjusted for and why they were included | 11 |
| | 16b | Report category boundaries when continuous variables were categorized | n/a |
| | 16c | If relevant, consider translating estimates of relative risk into absolute risk for a meaningful time period | n/a |
| | 16d | Report results of any adjustments for multiple comparisons | 11 |
| Other Analyses | 17a | Report other analyses done—e.g. analyses of subgroups and interactions, and sensitivity analyses | n/a |
| | 17b | If numerous genetic exposures (genetic variants) were examined, summarize results from all analyses undertaken | n/a |
| | 17c | If detailed results are available elsewhere, state how they can be accessed | n/a |
| **DISCUSSION** | | | |
| Key Results | 18 | Summarise key results with reference to study objectives | 17-18 |
| Limitations | 19 | Discuss limitations of the study, taking into account sources of potential bias or imprecision. Discuss both direction and magnitude of any potential bias | 18-20 |
| Interpretation | 20 | Give a cautious overall interpretation of results considering objectives, limitations, multiplicity of analyses, results from similar studies, and other relevant evidence | 20 |
| Generalisability | 21 | Discuss the generalisability (external validity) of the study results<br>Other information | 18-20 |
| **FUNDING** | | | |
| | 22 | Give the source of funding and the role of the funders for the present study and, if applicable, for the original study on which the present article is based | 21 |

*Give information separately for cases and controls in case-control studies and, if applicable, for exposed and unexposed groups in cohort and cross-sectional studies.

BRAIN A JOURNAL OF NEUROLOGY
ScholarOne, 375 Greenbrier Drive, Charlottesville, VA, 22901 Support (434) 964 4100
